# TcrDesign: De novo design of epitope specific full-length T cell receptors

**DOI:** 10.64898/2026.01.15.699824

**Authors:** Kaixuan Diao, Jing Chen, Xiangyu Zhao, Tao Wu, Die Qiu, Weiliang Wang, Haopeng Wang, Xue-Song Liu

**Author notes:** These authors contributed equally (Kaixuan Diao, Jing Chen).

## Abstract

T cell receptors (TCRs) are essential for adaptive immune recognition. Recently computational tools have been developed to predict the interactions between TCR and epitope, and artificial intelligence models have been proposed to generate the complementarity determining region 3 (CDR3) region of β chain TCR. However, de novo design of experimentally validated, functional, full-length epitope-specific TCRs remains a significant challenge. Here we developed TcrDesign, a deep learning framework based on large-scale unlabeled TCR and epitope datasets to generate epitope-specific, full-length TCRs. TcrDesign comprises two modules: TcrDesign-B for TCR-pMHC binding prediction with state-of-the-art accuracy, and TcrDesign-G for functional full-length TCR sequence generation. Pre-trained on large-scale unlabeled datasets using transformer-based architectures, TcrDesign achieves state-of-the-art performance in both TCR-epitope binding prediction and de novo TCR sequence generation. Furthermore, epitope-major histocompatibility complex (MHC) binding and functional activation of TcrDesign-generated TCRs were experimentally validated. TcrDesign provides an efficient and modular approach for designing epitope-specific full-length TCRs, with experimental validation confirming its utility.

## INTRODUCTION

T cell receptors (TCRs) play a critical role in the immune system by allowing T cells to recognize and respond to a diverse range of antigenic peptides presented by major histocompatibility complex (MHC) molecules. The interactions between TCRs and peptide-MHC complexes (pMHC) form a highly diverse and complex specific recognition system, which is crucial for the adaptive immune system (Dossa et al., 2018; Joglekar and Li, 2021; Krogsgaard and Davis, 2005). Although several experimental assays can determine specific TCR-epitope interactions, they are time-consuming and expensive (Bentzen et al., 2016; Dolton et al., 2015; Moravec et al., 2024; Zhang et al., 2021). The use of *in silico* methods to quickly and accurately determine and generate epitope-specific TCRs is an appealing alternative.

Nowadays, substantial TCR sequencing data have been accumulated, leading to the establishment of comprehensive TCR-related immune databases such as VDJdb, IEDB, McPAS-TCR, TCRdb, PIRD (Bagaev et al., 2020; Chen et al., 2021; Shugay et al., 2018; Tickotsky et al., 2017; Vita et al., 2019; Zhang et al., 2019). The prediction of TCR-pMHC interactions has been the primary focus of several recent studies, and recent advances in TCR specificity prediction primarily employ two approaches: physical distance quantification-based methods and deep learning methods (Croce et al., 2024; Dash et al., 2017; Jensen and Nielsen, 2023; Lu et al., 2021; Mayer-Blackwell et al., 2021; Springer et al., 2020; Springer et al., 2021). Previous studies have drawn several key conclusions regarding TCR-pMHC interaction predictions (Deng et al., 2023; Jensen and Nielsen, 2023; Meysman et al., 2023; Nielsen et al., 2024): firstly, accurate predictions require the use of paired α and β chains; secondly, the availability of sufficient TCR-pMHC interaction data is essential; and thirdly, extending predictions to novel epitopes (also known as unseen epitopes) without known TCRs remains challenging.

The ability to rapidly and efficiently generate specific TCR sequences for target antigens could significantly advance immunotherapy strategies such as TCR-T (Baulu et al., 2023; Brightman et al., 2020; Xu et al., 2022). Currently, despite advancements in TCR design, significant challenges persist in this field (Lin et al., 2024; Yang et al., 2023; Zhou et al., 2024b): (1) Limitations in generating full-length TCRs. Existing studies focus on generating epitope-specific βCDR3 regions rather than full-length TCRs, due to the abundance of βCDR3 data compared to the scarcity of αCDR3 data; (2) Lack of experimental validation. Although some tools theoretically demonstrate the capability to generate TCR sequences, these AI-generated TCR sequences have not been tested for functionality, which limits their reliability and feasibility in practical applications.

In this study, we develop TcrDesign, which leverages large language model technology to generate full-length TCR sequences for target epitopes and performs experimental verification, thereby addressing the aforementioned limitations. Our approach leverages large-scale, unlabeled TCR and epitope data to pre-train several language models, enabling the extraction of meaningful biological representations for downstream applications. TcrDesign comprises two modules: TcrDesign-B (the binding model) and TcrDesign-G (the generation model). TcrDesign-B accurately predicts interactions between epitopes and TCRs, achieving state-of-the-art performance. Meanwhile, TcrDesign-G demonstrates its effectiveness in generating diverse epitope-specific TCRs, which was subsequently validated through wet experiments.

## RESULTS

### TcrDesign workflow

As illustrated in **Figure 1A**, we designed and screened epitope-specific TCRs using the TcrDesign pipeline. The TcrDesign-G module was constructed to generate epitope-specific TCRs, which were then filtered and screened based on binding probability using the TcrDesign-B module. Structural modeling of the selected TCRs was performed with TCRmodel2 (Yin et al., 2023), and we chose TCRs with high structural confidence scores. Finally, we synthesized the candidate TCRs and conducted functional validation via experimental assays (see “Methods” for details).

**Figure 1.**
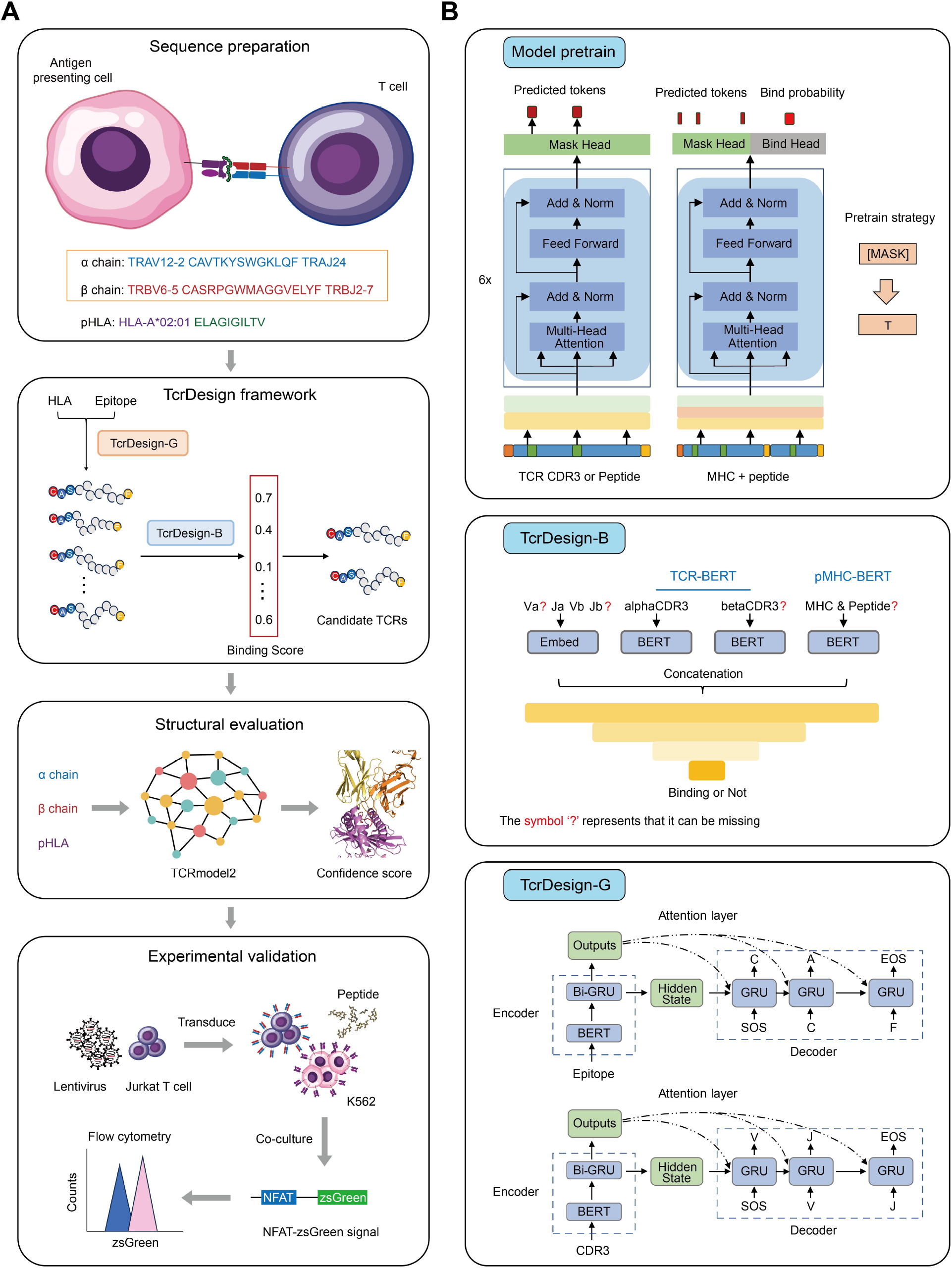
Generating full-length epitope-specific TCRs with TcrDesign. (**A**) Pipeline for the design and validation of full-length epitope-specific TCRs. TCRs are initially designed and screened for epitope specificity using the TcrDesign algorithm. These TCRs then undergo structural modeling, followed by further selection based on confidence scores from structural prediction tools. The selected TCRs are subsequently synthesized and validated for functional activity through in vitro experiments. (**B**) Overall framework of TcrDesign. BERT model architecture is employed for the pre-training on TCR and pMHC sequences to extract meaningful representations. The pre-training process for TCRs utilizes a MLM approach, while a dual-objective strategy is applied for pMHC, incorporating both masked recovery and the prediction of MHC and peptide binding. This results in the creation of TCR-BERT and pMHC-BERT pre-trained models for downstream applications. TcrDesign-B is developed based on these pre-trained models to predict TCR-pMHC binding, whereas TcrDesign-G is a TCR generation model that leverages attention mechanisms and a Seq2Seq architecture, specifically designed for generating full-length epitope-specific TCRs.

We gathered extensive public data from databases such as IEDB (Vita et al., 2019), VDJdb (Bagaev et al., 2020), MCPAS (Tickotsky et al., 2017), and TCRdb (Chen et al., 2021) to train several Bidirectional Encoder Representations from Transformers (BERT) models. The pre-training strategy employed was masked language modeling (MLM), which involved randomly masking 20% of the amino acids: 80% were replaced by a masking character, 10% by random characters, and 10% remained unchanged. This approach endowed the models with error correction capabilities and facilitated the generation of more meaningful representations (see “Methods”). We pre-trained three models: αCDR3-BERT, βCDR3-BERT, and pMHC-BERT, designed to extract representations for the TCR αCDR3 region, βCDR3 region, and pMHC complex, respectively. While αCDR3-BERT and βCDR3-BERT utilized a single masking pre-training task, pMHC-BERT incorporated an additional pMHC binding pre-training task to enhance feature extraction (**Figure 1B**). Using these pre-trained models, we established the TcrDesign-B module for predicting TCR-pMHC binding and the TcrDesign-G module for generating epitope-specific TCRs, thus forming the complete TcrDesign pipeline. Unlike previous models that focused solely on CDR3 sequence generation, our approach recognizes the critical importance of full-length TCR sequences. By leveraging V(D)J recombination principles, we developed a methodology utilizing pre-trained generative models to predict VJ gene segments from CDR3 sequences, thereby enabling full-length TCR sequence generation (**Figure 1B**).

### Evaluation of the performance of pre-trained models

TCRs are generated through VDJ rearrangement, with the CDR3 region at the VJ gene junction displaying remarkable diversity and extensive interactions with epitopes (Davis and Bjorkman, 1988; Krogsgaard and Davis, 2005). To capture the nuances of the TCR CDR3 region, we initially pre-trained TCR-BERT, which comprises two models: αCDR3-BERT and βCDR3-BERT (details in Methods and **Figure S1**). Given the limited data available for αCDR3 compared to βCDR3, we fine-tuned the pre-trained βCDR3-BERT model using the αCDR3 dataset, resulting in αCDR3-BERT. As shown in **Figure S2**, the fine-tuned αCDR3-BERT achieved superior accuracy in masked sequence recovery, indicating enhanced representation of the CDR3 region. Comparative analysis demonstrated that TCR-BERT achieved superior performance in masked amino acid recovery for both αCDR3 and βCDR3 regions when benchmarked against previously published models, suggesting enhanced learning of the underlying biological principles governing TCRs (**Figure 2A**).

**Figure 2.**
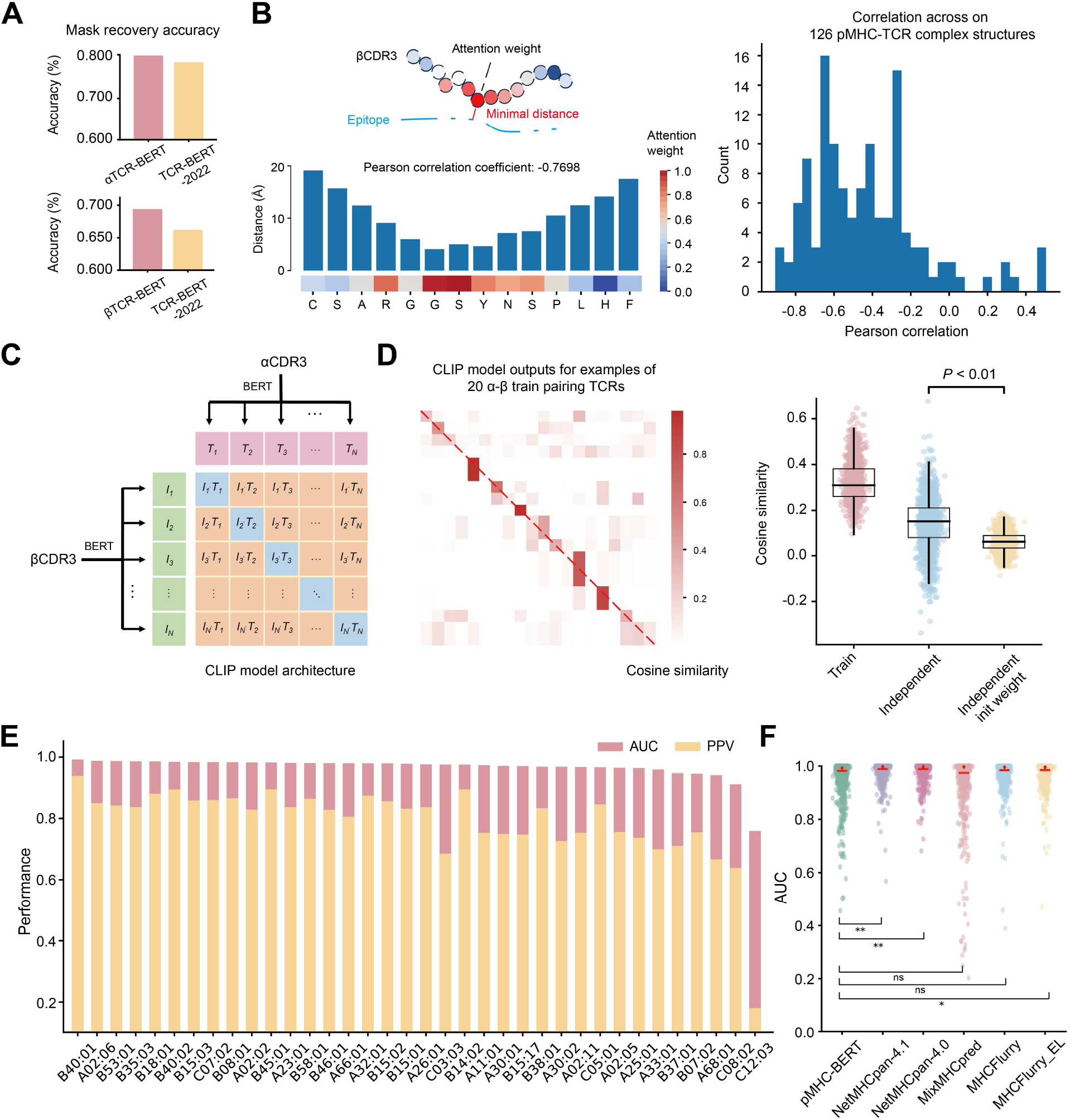
Pre-trained models efficiently learn the grammars within TCRs and pMHCs. (**A**) Performance comparison of αCDR3-BERT and βCDR3-BERT with TCR-BERT-2022, evaluated based on amino acid recovery accuracy. (**B**) Attention weight analysis highlights the importance of key amino acid residues in βCDR3 for antigen peptide binding. The left panel presents an example where warmer colors indicate higher attention weights. Pearson correlation coefficients were calculated between the Ca-Ca distances of βCDR3 amino acids and the epitope, as well as their corresponding attention weights. The right panel shows the distribution of Pearson correlation coefficients for 126 pMHC-TCR complexes. (**C**) Schematic representation of the CLIP model designed to learn the pairing relationships between TCR α and β chains. (**D**) The left panel displays model outputs for 20 training TCR pairs, while the right panel illustrates the model’s performance on an independent validation set after training. “Train” indicates performance on the training set, “Independent” refers to the independent validation set, and “Independent init(ial) weight” represents the performance of the untrained model on the independent validation set. A two-sided Student’s t test was applied. (**E**) Predictive performance of pMHC-BERT across various MHC subtypes in the CD8 EL SA dataset. (**F**) Comparison of the prediction performance of pMHC-BERT against other models on the CD8 epitope dataset.

We further analyzed the TCR features by examining the attention weights for the identification of key amino acid residues in βCDR3 (**Figure 2B**). Using a representative TCR βCDR3-epitope pair as a case study, we correlated the βCDR3 amino acid-level attention weight scores with their distances to the epitope. This analysis revealed that central amino acid residues of βCDR3 are critical for binding, with a correlation coefficient of -0.7698 (**Figure S3**).

Additionally, we collected 126 pMHC-TCR complex structures from the PDB (Sussman et al., 1998) and calculated correlation coefficients in the same manner. Our findings showed strong correlation between attention weights and physical distances, with correlation coefficients predominantly ranging from -0.4 to -0.8 (**Figure 2B**). This suggests that TCR-BERT implicitly captures the functional importance of individual amino acids in CDR3.

Previous studies have indicated an intricate pairing relationship between TCR α and β chains, particularly the patterns of V and J gene pairing (Carter et al., 2019; Grigaityte et al., 2017; Tanno et al., 2020). The Contrastive Language-Image Pretraining (CLIP) model, originally developed by OpenAI for image-text pairing, has demonstrated strong zero-shot learning capabilities (Radford et al., 2021). In this study, we leveraged the CLIP model architecture to explore the pairing relationship between TCR α and β chains. We fine-tuned the CLIP model using the pre-trained TCR-BERT as a foundation (**Figure 2C**). After training, we observed increased cosine similarity of features between paired TCR α and β chains compared to the untrained model (**Figure 2D**). This finding suggests that the model successfully learned the pairing relationship between TCR α and β chains.

To assess the pre-training effectiveness of our pMHC-BERT model (details in Methods and **Figure S1**), we conducted a comprehensive evaluation focusing on its peptide-MHC binding prediction capabilities. The evaluation utilized two independent validation datasets derived from the netMHCpan v4.1 method (Reynisson et al., 2020): the CD8 epitope dataset and the CD8 EL SA dataset (“Methods”). On the CD8 EL SA dataset, pMHC-BERT exhibited strong performance, achieving AUC values exceeding 0.9 for various MHC subtypes (**Figure 2E**). pMHC-BERT also demonstrated comparable performance when compared to other advanced pMHC binding prediction models (Gfeller et al., 2023; O’Donnell et al., 2020; Reynisson et al., 2020) on the CD8 epitope dataset (**Figure 2F and S4**). The observed MHC subtype-dependent performance variations in pMHC-BERT (**Figure S5**) are addressed comprehensively in the discussion section. These results demonstrate the effectiveness of pMHC-BERT pre-training.

### TcrDesign-B accurately predicts the interactions between TCRs and epitopes

Previous studies have shown that existing TCR binding prediction models fail to generalize to unseen epitopes, exhibiting low accuracy for epitopes not included in training sets while performing well on seen epitopes with multiple associated binding TCRs (Meysman et al., 2023; Moris et al., 2021; Nielsen et al., 2024). We built upon the pre-trained TCR-BERT and pMHC-BERT models to develop TcrDesign-B, which aims to improve generalization to unseen epitopes by leveraging large-scale unlabeled data.

We build two versions of TcrDesign-B: the pan-epitope version, which is trained on all available epitopes and TCRs, and the epitope-specific version, which consists of separate models for each individual epitope. In line with previous findings (Croce et al., 2024), we focused on epitopes with more than 40 associated binding TCRs for training, leading to the development of 39 epitope-specific TcrDesign-B models (**Table S1**). The performance of TcrDesign-B was evaluated using four additional datasets: 1) a held-out validation dataset, 2) an independent common epitope dataset, 3) an independent novel epitope dataset, 4) an independent IMMREP2023 dataset (Nielsen et al., 2024) (see “Methods”). We first trained the pan-epitope TcrDesign-B, which performed well on the validation dataset (**Figure S6A**). Next, we found that the epitope-specific TcrDesign-B significantly outperformed the pan-epitope version on the 39 epitopes that both models could predict (**Figure 3A**), a trend that was also evident on the common epitope dataset (**Figure S6B**). These findings indicate that training TcrDesign-B with all available TCR-epitope pairs is less effective than developing epitope-specific models, aligning with previous research (Croce et al., 2024; Jensen and Nielsen, 2023; Montemurro et al., 2022). As a result, we adopted a hybrid approach for the subsequent TcrDesign-B, employing epitope-specific models for the 39 epitopes and the pan-epitope model for all other epitopes. This strategy strikes a balance between predictive accuracy and broad applicability.

**Figure 3.**
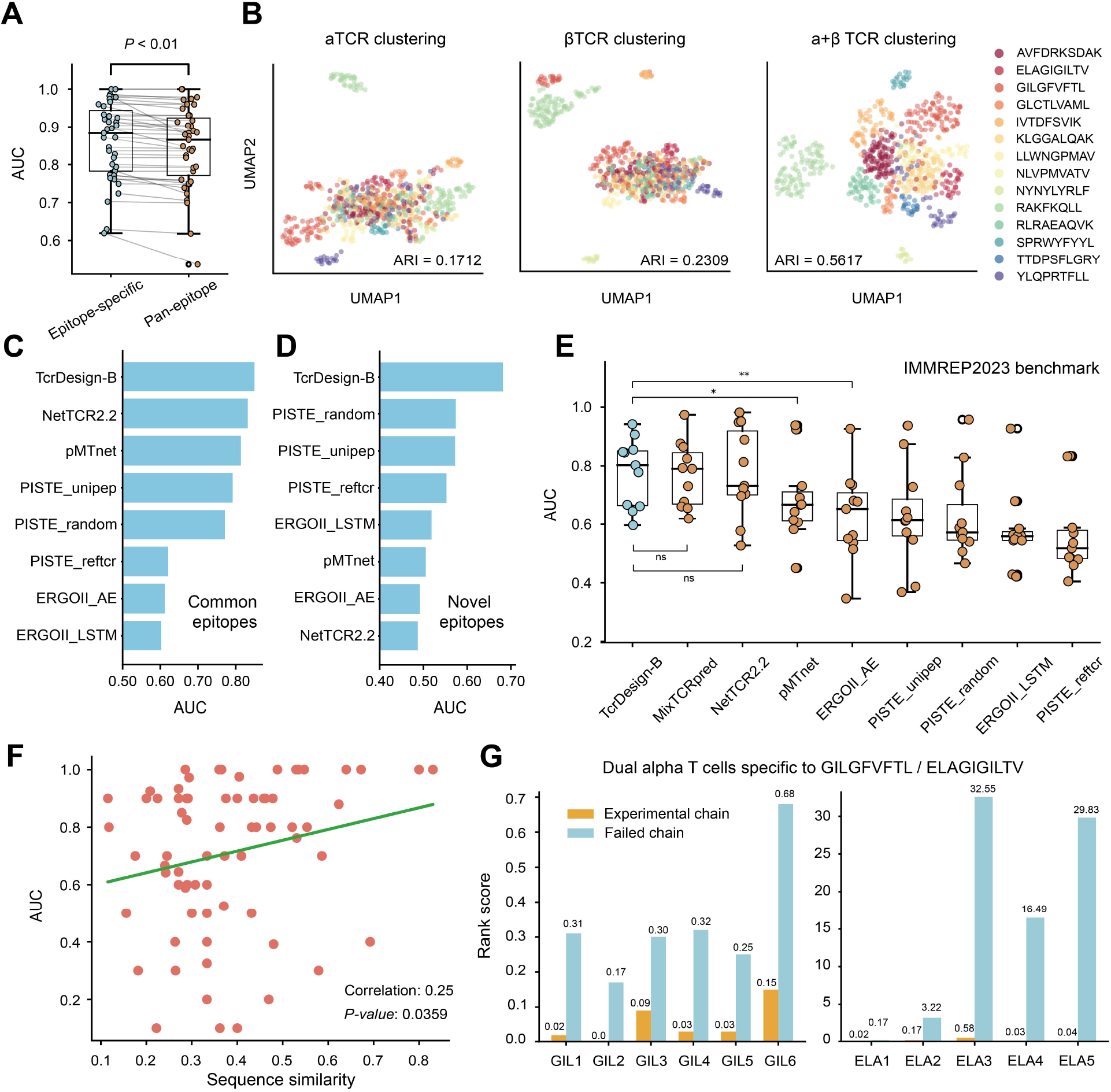
TcrDesign-B accurately predicts the interactions between TCRs and epitopes. (**A**) Performance comparison between the epitope-specific and pan-epitope versions of TcrDesign-B on the validation dataset, with each point representing an individual epitope. A two-sided paired Student’s t-test was conducted to assess significance. (**B**) UMAP dimensionality reduction of epitope-specific TCRs from the common epitope dataset utilizing TCR embedding extracted by TcrDesign-B. The ARI was employed to evaluate clustering performance, where higher values indicate better clustering. (**C**) Performance comparison of TcrDesign-B with other models on the common epitope dataset, where “common epitope” refer to those included in the training set. (**D**) Performance comparison of TcrDesign-B with other models on the novel epitope dataset, which includes epitopes not present in the training set. (**E**) Performance comparison of TcrDesign-B with other models on the independent validation dataset IMMREP2023, a paired two-sided t-test were applied. (**F**) Scatter plot illustrating the sequence similarity of epitopes in the novel epitope dataset compared to the most similar epitopes in the training set, correlated with TcrDesign-B performance as measured by AUC. Correlation and linear regression analysis, along with significance, are reported. (**G**) TcrDesign-B rank scores for two TCRs in dual alpha T cells specific to GIL and ELA epitopes. “Experimental chains” refer to TCRs capable of binding GIL or ELA to elicit T cell responses, while “failed chains” are those that do not elicit T cell responses.

The absence of TCR information (TCR α or β chain) profoundly impacts on model performance, as supported by previous studies (Fast et al., 2023; Montemurro et al., 2022; Springer et al., 2021). In this study, we examined how TCR information affects TcrDesign-B’s performance using a clustering approach. We selected 14 epitopes from the common epitope dataset and 10 from the novel epitope dataset. Each epitope has at least two associated binding TCRs. TCR embedding was extracted from TcrDesign-B, and we used UMAP (Uniform Manifold Approximation and Projection) for dimensionality reduction. The adjusted rand index (ARI) was employed to assess the quality of the clustering (see “Methods”). Our analysis showed that clustering based solely on either TCR α or β chain information was insufficient in both the common epitope dataset (**Figure 3B**) and the novel epitope dataset (**Figure S7**). However, clustering that incorporated all available TCR chain information yielded significantly improved results. These findings indicate that both TCR α and β chain information are essential for TcrDesign-B, the absence of either one markedly diminishes model performance.

To provide a clearer evaluation of TcrDesign-B’s performance, we conducted a fair comparison against other TCR binding prediction models. All models performed well in both the common epitope dataset and the IMMREP2023 dataset (**Figure 3C and E**), demonstrating that they can learn TCR binding rules from adequate training data. Both TcrDesign-B and MixTCRpred exhibited superior performance (**Figure 3E and S8**). However, in the novel epitope dataset comprising unseen epitopes, all models experienced a decline in predictive performance, with TcrDesign-B achieving the best performance (**Figure 3D**). This finding suggests that large-scale and unsupervised pre-training effectively captures biological knowledge that is advantageous for TCR binding prediction.

Despite the general inaccuracy of TCR binding prediction models for unseen (novel) epitopes, we found that the predictive performance significantly improves as the sequence similarity between unseen epitopes in the novel epitope dataset and seen epitopes in the training set increases (**Figure 3F**, see “Methods”). This suggests that when TcrDesign-B is applied to a new epitope, the presence of highly similar epitopes in the training set—particularly those with over 40 associated TCRs—can enhance prediction accuracy. These findings align with previous findings (Deng et al., 2023; Grazioli et al., 2022; Hudson et al., 2023). Additionally, it is important to note that most T cells typically express a single α and β chain to form a unique TCR (Brady et al., 2010). However, prior research indicates that approximately 10% of T cells express two different α chains, while double β chains are present in less than 1% of T cells, suggesting that dual α chain T cells are relatively common (Han et al., 2014; Heath et al., 1995; Padovan et al., 1993; Schuldt and Binstadt, 2019). Using TcrDesign-B, we validated the previously experimentally confirmed dual α chain T cells specific to HLA-A*02:01/GILGFVFTL and HLA-A*02:01/ELAGIGILTV pMHCs, demonstrating that TcrDesign-B can effectively identify epitope-specific TCR chains (**Figure 3G and Table S2**).

We extended TcrDesign-B by adding a regression head and fine-tuning it to predict amino acid-level interactions between βCDR3s and epitopes. The fine-tuning dataset consists of 110 TCR-pMHC complex structures obtained from the Protein Data Bank (PDB) database, while validation was conducted using a test set of 11 samples. Two examples (PDB IDs: 7QPJ and 8GON) illustrate that the fine-tuned model effectively captures amino acid-level interactions, with strong positive correlation between predicted and actual contact scores (normalized Ca-Ca distances) (**Figure S9**).

### TcrDesign-G enables generation of epitope-specific full-length TCRs

Generating epitope-specific full-length TCR sequences remains a significant challenge, and existing methods, such as GRATCR (Zhou et al., 2024b), can only produce epitope-specific βCDR3 sequences, which do not fully meet practical requirements. To address this challenge, we developed the TcrDesign-G model by leveraging pre-trained knowledge to generate epitope-specific full-length TCR sequences. The model utilizes the V(D)J recombination principle and a pre-trained generative architecture to predict VJ genes from CDR3 sequences. Using available training samples, we categorized epitopes as common (≥400 TCRs), rare (1-399 TCRs), or novel (0 TCR). Representative epitopes from each category were selected to thoroughly assess the performance of TcrDesign-G.

TcrDesign-G and GRATCR were directly compared on common epitopes, which are present in both training data sets, to ensure fairness. We used the BLOSUM62 score to evaluate the functional conservation of the TCRs generated by our model. BLOSUM substitution matrices are derived from evolutionary relationships among proteins to indicate functional similarity (Eddy, 2004). Protein functional similarity can be quantified by calculating BLOSUM62 scores of protein sequences (see “Methods”). TcrDesign-G exhibited high functional similarity to natural CDR3s across diverse epitopes, underscoring the functional validity of these generated sequences (**Figure 4A, 4B and S12**). TcrDesign-G outperformed GRATCR (**Figure 4A**) and significantly exceeded a random baseline model (starting with C, ending with F, and containing random amino acids in between). Regarding resource usage, TcrDesign and GRATCR exhibit similar runtime and GPU memory consumption, indicating comparable practicality for both in real-world applications (**Figure S10**). Furthermore, TcrDesign-G achieved robust performance on the binding score metric as calculated by TcrDesign-B, highlighting its effectiveness in generating epitope-specific TCRs. To facilitate a comparative evaluation of TcrDesign-G (a BERT-based generative model) and decoder-only architectures for TCR design, we trained a GPT baseline model. When evaluated on 12 representative epitopes, TcrDesign-G demonstrated superior performance (**Figure S11**), as discussed further in the Discussion.

**Figure 4.**
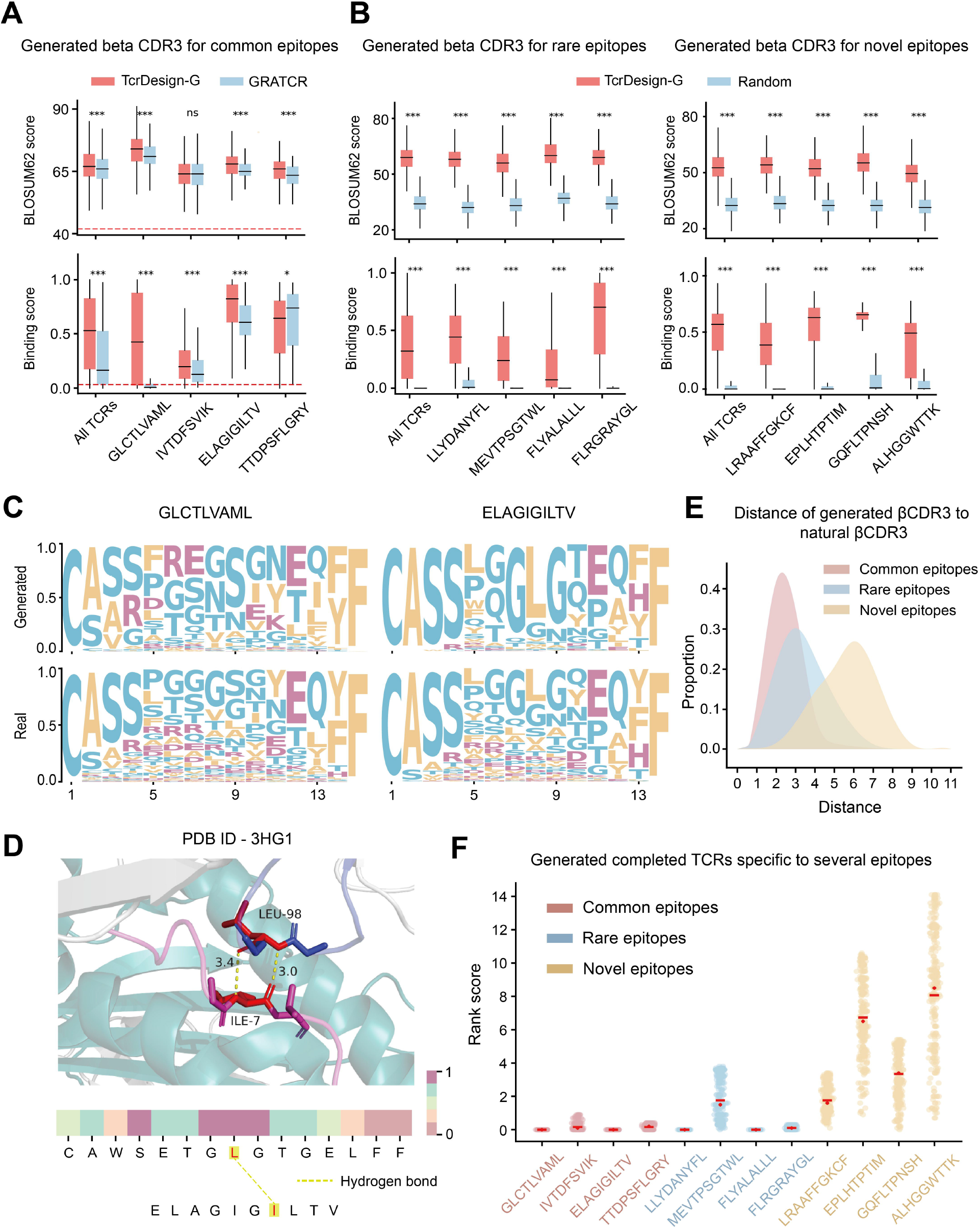
TcrDesign-G successfully generates epitope-specific TCRs. The performance of TcrDesign-G is evaluated using beta chain information for common epitopes (**A**) and rare or novel (unseen) epitopes (**B**). The BLOSUM62 score indicates the biological functional similarity between generated and natural sequences, while the binding score reflects the predicted binding probability as assessed by TcrDesign-B. Two-sided Student’s t tests were applied (*p<0.05, **p<0.01, ***p<0.001, “ns” denotes “not significance”). (**C**) Motif analysis demonstrates the generation of 1,000 βCDR3 sequences for the common epitopes GLCTLVAML and ELAGIGILTV, comparing these to the natural binding βCDR3 sequences. (**D**) Interaction analysis between the “GLG” motif within the ELAGIGILTV-HLA-A*02:01-specific TCR and ELAGIGILTV epitope (PDB ID 3HGI). The residues highlighted in red indicate those involved in the interaction, with the color reflecting the attention weights derived from TCR-BERT. (**E**) The density plot illustrates the distribution of the minimum edit distances between 10,000 generated βCDR3 sequences for common, rare, and novel epitopes and their corresponding natural βCDR3 sequences. (**F**) TcrDesign-G generated 10,000 full-length TCRs for common, rare and novel epitopes, respectively. The distributions of their TcrDesign-B rank scores are reported.

To assess the model’s capability to learn TCR syntax, we conducted a comparison between TcrDesign-G-generated TCRs and natural TCRs. The βCDR3 sequences generated by TcrDesign-G for two common epitopes captured important motifs (**Figure 4C**). The generated βCDR3s for ELAGIGILTV were particularly noteworthy, with the presence of the GLG motif (**Figure 4D**). However, such motifs were absent in CDR3s generated for novel epitopes (**Figure S13**). A similar phenomenon was observed in the generation of αCDR3 sequences (**Figure S14**). We further analyzed the TcrDesign-G-generated sequences. A typical 15-mer CDR3 sequence consists of two conserved segments (approximately the first 4 and last 4 amino acids of βCDR3, and the first or last 3 of αCDR3) and a highly variable middle region of about 7 amino acids. As anticipated, we observed an increase in the distance of CDR3 sequences generated by TcrDesign-G when transitioning from common to novel epitopes (**Figure 4E and S15**). For common epitopes, while the majority of generated sequences differed from known sequences by 2-4 amino acids, some differed by more than 5 amino acids. This finding suggests that TcrDesign-G can generate novel sequences for well-represented common epitopes rather than simply memorizing existing patterns. Further analysis of the length and amino acid distribution of the generated TCRs revealed that those produced by TcrDesign closely matched the native distribution (**Figure S16**). Finally, we generated full-length epitope-specific TCRs using TcrDesign-G and calculated rank scores using TcrDesign-B (see “Methods”), for common or rare epitopes, TcrDesign can generate top-ranked TCRs; however, its efficacy is constrained when dealing with novel epitopes, underscoring the persistent challenges in generalizing to unseen epitopes (**Figure 4F**).

Given that V(D)J recombination enables the inference of V and J genes from CDR3 sequences (Malu et al., 2012; Schatz and Ji, 2011; Van Gent et al., 1996), TcrDesign-G can be utilized to generate full-length TCRs. We implemented a model (**Figure 1B**, see “Methods”) to infer VJ genes from CDR3 sequences. Our analysis revealed that although J genes could be accurately predicted from CDR3 sequences, the identification of V genes demonstrated lower precision. However, accuracy significantly improves when considering the top 5 predictions (i.e., the top 5 predicted VJ gene pairs from a CDR3 sequence; **Figure S17**). We hypothesize that the conserved nature of CDR3 start sequences, such as CASS or CAV, contributes to the difficulty in reliably inferring V genes. This analysis not only demonstrates the capabilities of TcrDesign-G but also highlights areas for future improvement, particularly in enhancing the model’s ability to generalize to novel epitopes and in refining the accuracy of V gene inference from CDR3 sequences.

### TCRs designed by TcrDesign show functional activation

We generated full-length TCRs for two common epitopes and visualized their characteristics through dimensionality reduction. The distinct clustering of TCRs for different epitopes clearly demonstrated the epitope specificity of TcrDesign-generated TCRs (**Figure 5A**). To further evaluate TcrDesign’s performance, we generated nine full-length TCRs for the common epitope ELAGIGILTV and HLA-A*02:01 (**Table S3**). Nine candidate TCRs were transduced into reporter Jurkat T cell lines via lentivirus to characterize the pMHC tetramer binding specificity and TCR-mediated immune response. A previously reported TCR was used as a positive control (Harris et al., 2016), while endogenous TCR from wild-type Jurkat cells acted as a negative control. The results demonstrated that three TcrDesign-generated TCRs (ELA3, ELA4 and ELA6) exhibited specific binding to the HLA-A*02:01-ELAGIGILTV tetramer (**Figure 5B**). Furthermore, these candidate TCRs mediated antigen-specific responses against ELAGIGILTV-pulsed T2 cells, as demonstrated by CD69 upregulation, NFAT-ZsGreen reporter activation (**Figure 5C, 5D and S19)** and IL-2 secretion in co-culture assays **(Figure 5E)**. We generated TCR-T cells by transducing lentivirus carrying the positive, ELA3, ELA4 and ELA6 TCR genes to primary T cells. Bioluminescence-based cytotoxicity assay revealed substantial killing of ELAGIGILTV-pulsed T2 cells by ELA3, ELA4 and ELA6 TCR-T cells (**Figure 5F and S20**). To offer mechanistic insight, we have included computational analyses of the binding energy and docking scores for the generated TCR–pMHC complexes. The results indicate that the three functionally validated TCRs (3, 4, and 6) ranked among the top candidates, suggesting superior complex stability (**Figure S21**). We also confirmed that the baseline luminescence of T2-luciferase cells was unaffected by 1 μg/mL peptide incubation (16 h), ruling out potential artifacts from differential cell viability (**Figure S22**). These functional assay results confirm TcrDesign’s ability to generate functional TCRs, highlighting its potential applications in immunotherapy design. We further identified the natural TCRs most similar to our designed sequences to analyze the relationship between sequence divergence and experimental performance. We observed that our designed TCRs exhibit distinct divergence from known natural sequences, in both V, J gene usage and CDR3 composition.

**Figure 5.**
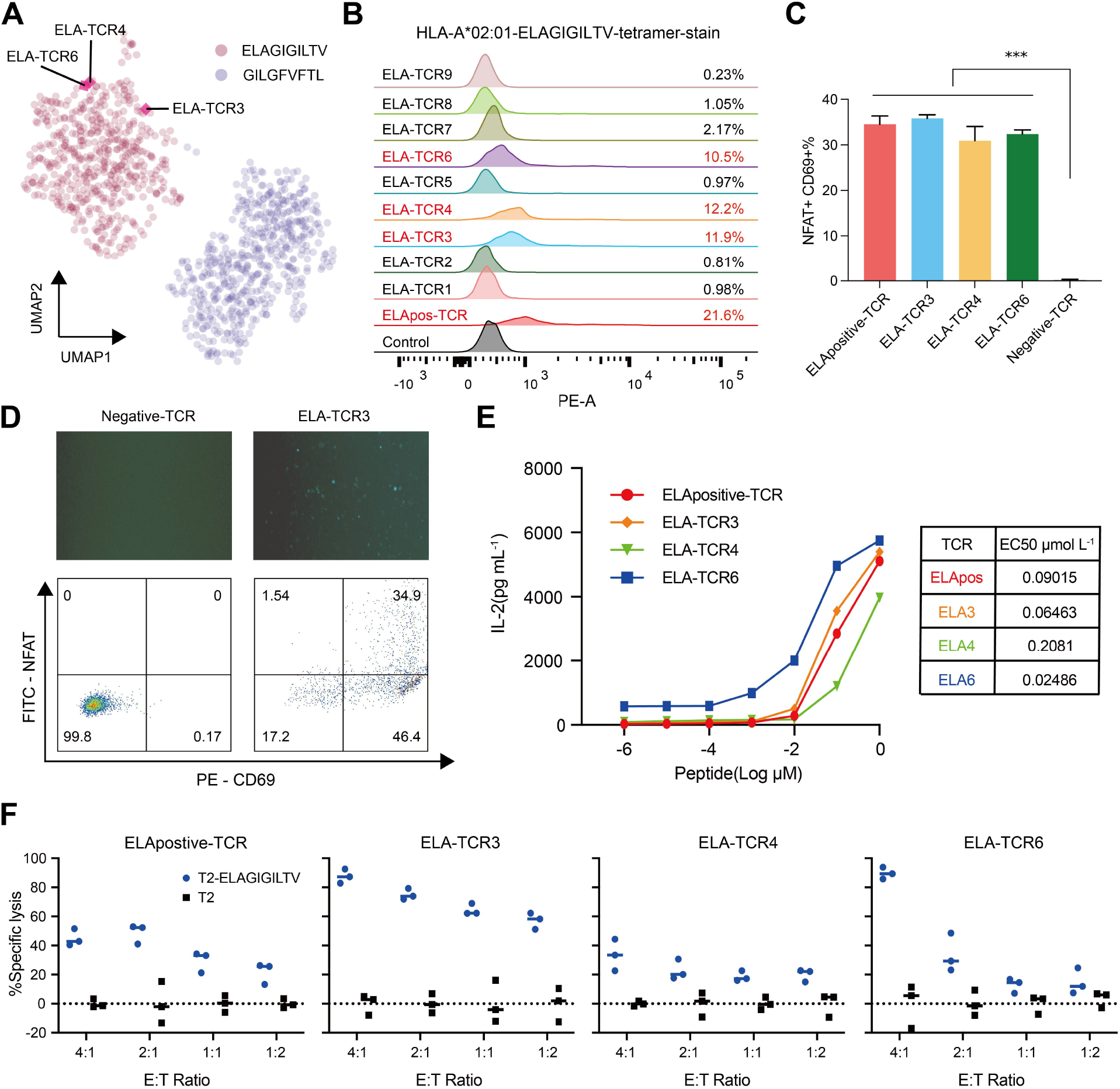
TCRs designed with TcrDesign show functional activation. (**A**) UMAP dimensionality reduction was performed on generated TCRs for two distinct epitopes, utilizing TCR embedding extracted by TcrDesign-B. (**B**) Binding assay of HLA-A*02:01-ELAGIGILTV tetramer to the nine generated TCRs expressing Jurkat cells was analyzed by flow cytometry. Three candidate TCRs ELA-TCR3, ELA-TCR4 and ELA-TCR6 exhibited specific binding to the tetramer. (**C**) Activation markers NFAT and CD69 levels of reporter Jurkat NFAT-ZsGreen cells expressing candidate TCRs ELA-TCR3, ELA-TCR4, ELA-TCR6 and ELApositive TCR after incubation with ELAGIGILTV-pulsed T2 cell for 24 hours, each sample was tested in triplicate. (**D**) The NFAT-ZsGreen signal and CD69 level of ELApositive-TCR and ELA-TCR3 were determined by fluorescence imaging and flow cytometry analysis. (**E**) Engineered Jurkat T cells were co-cultured with T2 cells and serially diluted peptides for 24 h. Co-cultured supernatants were analyzed by ELISA for secreted IL-2. (**F**) TCR-T cells at various E:T ratios were co-cultured with luciferase-transduced T2 cells. The % specific lysis of T2-luciferase (black) and 1ug/mL ELAGIGILTV pulsed T2-luciferase cells (blue) obtained by bioluminescence assay is plotted against multiple E:T ratios. Dots in the figure represents three replicates.

Importantly, proximity to natural TCRs did not predict experimental success (**Figure S23**). For instance, ELA-TCR1, the design most similar to natural sequences, showed no reactivity, whereas ELA-TCR6, which exhibits significant divergence, demonstrated a robust response. These results indicate that the model can generalize beyond the natural repertoire by learning underlying biological constraints rather than merely inferring from known sequences.

To comprehensively assess potential off-target effects, we performed additional experimental validations using both a panel of human cancer cell lines (including HLA-A*02:01-homozygous AGS cells, non-HLA-A*02:01 A549 cells, and 293T cells) representing diverse self-peptide backgrounds and alanine scanning of the target peptide. TCR activation was quantified via NFAT-luciferase reporter assays measuring NFAT+CD69+ responses. The results indicated negligible off-target reactivity against self-peptides across all tested cell lines. However, in alanine scanning, all TCRs exhibited cross-reactivity toward peptides with flanking residue substitutions, while responses to central mutations were minimal. This observation is consistent with the structural disruption when mutations happen at core binding residues (**Figure S24 and S25**). These results suggest that the generated TCRs may exhibit off-target effects in vivo. Consequently, future generative models should incorporate strategies for identifying and filtering TCRs with potential cross-reactivity. Furthermore, additional validation, including studies in animal models, will be essential to thoroughly assess the cross-reactivity of the designed TCRs.

## DISCUSSION

We present TcrDesign, an advanced deep learning algorithm for generating epitope-specific TCRs, representing a breakthrough in full-length TCR generation for practical applications. By leveraging pre-training on large-scale unlabeled TCR and pMHC datasets, TcrDesign achieves state-of-the-art performance in both TCR binding prediction and generation tasks. Furthermore, we successfully designed TCRs for the common cancer-related epitope ELAGIGILTV using TcrDesign, with generated TCRs effectively binding to the epitope and activating T cells. The computational and experimental results highlight TcrDesign’s potential impact in the field of TCR design. Additionally, the traditional single-cell-based TCR discovery process, although effective, suffers from low throughput, high experimental difficulty, and the potential inability to identify optimal epitope-specific TCRs, rendering it time-consuming and labor-intensive (Wang et al., 2025; Xiong et al., 2022). TcrDesign can rapidly generate hypotheses for diverse epitopes, which substantially narrows the search space for possible sequences and enhances the efficiency of TCR discovery.

The development of TcrDesign involved a deliberate selection of a BERT-based encoder architecture, a decision motivated by the specific challenges and objectives of TCR design. While large, decoder-only language models have demonstrated exceptional capabilities in broad sequence generation tasks, our goal was to construct a framework that prioritizes high-fidelity feature extraction from TCRs and generation of structurally coherent, full-length TCRs (Xu et al., 2026). The bidirectional context understanding inherent to Transformer encoders is particularly advantageous for modeling the complex, non-linear interactions within TCR-pMHC complexes, which are crucial for predicting TCR binding. Well-established BERT models frequently demonstrate superior or at least comparable performance to decoder-only models, particularly when fine-tuned for specific domain tasks, owing to their bidirectional architecture (Benayas et al., 2024; Raffel et al., 2023). Our results also show that the BERT-based generation model has an advantage in the current TCR design task (**Figure S11**).

The observed MHC subtype-dependent performance disparities in pMHC-BERT may be attributed to limited ligand datasets for certain MHC molecules (e.g., HLA-C and HLA-DQ). Similar phenomena have been reported in NetMHCpan, where MHC molecules with scarce data exhibit lower predictive accuracy (Reynisson et al., 2020). This may be due to difficulties in obtaining experimental data for different MHC molecules, resulting in varying amounts of training data. These insights suggest that data scarcity is a primary driver of the disparities, though contributions from biological and algorithmic elements warrant further exploration in future work with expanded datasets for underrepresented alleles.

Despite advancements in TcrDesign, limitations remain in TCR generation. Benchmark studies show that TCR models often underperform with unseen epitopes (Meysman et al., 2023; Nielsen et al., 2024). While TcrDesign shows improvement in handling unseen epitopes, its ability to generate TCRs for these epitopes remains suboptimal. The incorporation of structural information has been demonstrated to enhance model generalization in multiple studies (Su et al., 2023; Sun et al., 2025; Zhang et al., 2025). Although experimentally determined TCR-pMHC structures remain scarce, a recent investigation has performed comparative and systematic evaluations of tools for predicting TCR structures and TCR-pMHC complex structures, highlighting the effectiveness of TCRmodel2 and AlphaFold2/3 in modeling TCR-pMHC complexes (Shi et al., 2025). Leveraging the reliable TCR-pMHC structures generated by these tools, which provide more comprehensive information than sequence data alone, has been valuable for improving model performance, especially with limited training samples. Therefore, future work may benefit from integrating structural information to address current limitations in generating TCRs for novel epitopes. It is important to bear in mind that TCR structural modeling remains challenging, primarily due to the difficulty in accurately predicting the conformation of CDR loops. With continued advances in deep learning algorithms and structural biology, TCR–pMHC modeling is expected to improve significantly, which will greatly facilitate and accelerate the development of TCR design.

An additional limitation of this study is the lack of consideration for thymic selection, a fundamental process in shaping the natural TCR repertoire. Thymic selection ensures that T cells are capable of recognizing foreign antigens effectively while maintaining tolerance to self-antigens, thereby upholding immune homeostasis. Disruption of such regulatory mechanisms has been shown to contribute to autoimmune pathology (Shao et al., 2025). Notably, current AI-driven TCR design methodologies, including TcrDesign, overlook this critical tolerance checkpoint. To enhance the biological relevance and safety of AI-generated TCRs, future efforts could incorporate modeling of thymic selection pressures - for instance, by adopting frameworks similar to those proposed by Isacchini and colleagues (Isacchini et al., 2021) - to effectively filter and prioritize candidate TCRs.

In our experiments, validation was limited to the cellular level for designs targeting the common tumor antigen ELAGIGILTV, offering valuable yet limited insight into T-cell activation and cytotoxicity. The broader TCR design field remains challenged by difficulties in generalizing to unseen peptides, and designing TCRs specific to neoantigens for use in complex clinical settings presents formidable obstacles. Lessons from antibody design suggest that success rates for novel epitopes may be below 1% (Bennett et al., 2025), underscoring the need for more diverse experimental and clinical data to advance TCR design. Furthermore, TCR cross-reactivity represents a critical factor for clinical translation. Comprehensive filtering strategies against self-peptide databases will be needed in future iterations to address autoimmunity risks. Although our initial designs for ELAGIGILTV showed no detectable off-target effects against self-peptides, off-target binding was observed in alanine scanning assays. This finding prompted us to propose that additional validation, including studies in animal models, will be essential to assess the cross-reactivity effects of the designed TCRs. Broader experimental assessments encompassing multiple epitopes and more comprehensive cytotoxicity assays will be needed to fully evaluate the functionality of designed TCRs. Moreover, training on larger real-world datasets and developing more innovative model architectures will enhance the generalization capacity of TcrDesign and its utility in addressing practical challenges.

It should also be noted that in clinical applications of TcrDesign-generated TCRs for TCR-T therapy, in addition to off-target effects, consideration must be given to the potential mispairing between introduced and endogenous TCR chains—an issue that has been extensively studied and addressed through various engineering strategies (Legut et al., 2018; Su and Liu, 2025). Although TCR-T cells generally exhibit less severe cytokine release syndrome (CRS) than CAR-T cells, the risk of serious CRS remains and requires proactive clinical management. Fortunately, several approaches have been developed to effectively mitigate CRS (Leclercq et al., 2022; Rodrigues Dos Santos et al., 2024). AI-driven TCR design represents only the initial step; further engineering modifications are essential to optimize the functionality and safety of these designed TCRs for therapeutic use.

TCRs generally have lower binding affinities to pMHC compared with the binding between antibody and antigen, and TCR-pMHC binding does not always lead to T-cell activation (Spindler et al., 2020; Stone and Kranz, 2013). In general, natural TCRs exhibit a specific relationship between affinity and activity: when affinity is greater than 1-10 μM, affinity and activity are positively correlated; however, when affinity is less than 1-10 μM, further enhancements in affinity do not lead to corresponding increases in activity (Stone and Kranz, 2013; Zhong et al., 2013). The positive control TCR used in this study possesses high affinity (Kd = 0.129 μM), yet its activity is comparable to those of our generated TCRs (using tetramer binding experiments as a proxy for affinity and EC50 values as a measure of activity). This finding is consistent with prior research literature, indicating that the TCRs we generated display affinity-activity relationships akin to those observed in natural T cells.

In summary, TcrDesign serves as a valuable tool for TCR design, facilitating rapid TCR design through (i) quick TCR-epitope binding prediction and (ii) full-length epitope-specific TCR generation. We anticipate that with more data, TcrDesign’s capabilities will continue to be improved, offering valuable insights for TCR research.

## METHODS

### Datasets

#### pMHC presentation datasets

To prepare pMHC data for pre-training, we collected and filtered linear peptides ranging from 8 to 21 amino acids in length from IEDB database (Vita et al., 2019), ensuring that all peptides were derived from human sources (Type I peptides of 8-15 amino acids and Type II peptides of 13-21 amino acids). To enhance our dataset, we also included training data from netMHCpan v.4.1 (Reynisson et al., 2020) and netMHCIIpan v.4.3 (Nilsson et al., 2023). This consolidation resulted in a final dataset comprising 7,552,312 pMHC pairs across 306 HLA alleles. For binding affinity (BA) data, we implemented a specific scaling and normalization formula to facilitate integration with eluted ligand (EL) data for model training.

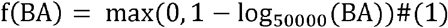

The upper limit for BA is generally set at 50,000, which ensures that *f*(*BA*) stays within the range of 0 to 1. We randomly selected 1% of the data to form a validation set, while the remaining data was used for model training. The training dataset comprises 7,476,788 samples, whereas the validation dataset contains 75,524 samples, resulting in a positive-to-negative sample ratio of approximately 1:9. To assess the pMHC-BERT’s predictive performance, we utilized the CD8 epitope dataset and the CD8 EL SA dataset from NetMHCpan v4.1 as benchmarks.

#### Pre-training peptide and TCR datasets

We extracted peptides from the pMHC presentation dataset, creating a deduplicated dataset of 6,539,012 unique peptides. The pre-training TCR dataset specifically focused on the critical CDR3 region, with separate training for the α and β chains to optimize performance. We gathered βCDR3 data from TCRdb (Chen et al., 2021), ImmuneCODE (Dines et al., 2020; Nolan et al., 2020), and other relevant literature (Carter et al., 2019; Tanno et al., 2020), yielding 40,339,559 unique entries. For αCDR3, we collected data from VDJdb (Bagaev et al., 2020) and related studies (Carter et al., 2019; Tanno et al., 2020), yielding 128,732 unique entries after deduplication.

#### TCR-pMHC binding datasets

The TCR-pMHC pairing data was gathered from various public datasets, including VDJdb, IEDB, McPAS (Tickotsky et al., 2017), the 10x Genomics dataset (Zhang et al., 2021), and MixTCRpred (Croce et al., 2024). To ensure data quality, we required that at least one chain (either alpha or beta) contain complete information, including both VJ genes and CDR3 sequences. CDR3 sequence lengths were restricted to 8-20 amino acids while preserving classical characteristics (starting with C and ending with F or W). After merging and filtering, we obtained a final dataset of 97,553 TCR-pMHC pairs for training and 5,894 for validation (95:5 split).

In addition, we collected data from relevant literature (Fast et al., 2023; Zhou et al., 2024a) to create two distinct benchmark datasets: a common epitope dataset, which includes peptides present in the training set, and a novel epitope dataset, consisting of peptides not found in the training set. After deduplication with the training set, the common epitope dataset was found to contain 704 positive pairs, while the novel epitope dataset included 112 positive pairs. Additionally, the dataset from the IMMREP2023 (Nielsen et al., 2024) competition was compiled as another benchmark dataset comprising 598 positive and 2886 negative pairs.

### Negative data sampling method

Building on previous research (Croce et al., 2024; Fast et al., 2023; Moris et al., 2021), we implemented several strategies to generate negative samples aimed at increasing diversity and enhancing the robustness of model predictions. Our specific steps were as follows:

1. Data augmentation: We expanded the positive training samples fivefold to mitigate data imbalance. For instance, using a TCR-pMHC paired sample (Va-CDR3a-Ja-Vb-CDR3b-Vb, peptide-MHC), we masked certain elements of the data. This included masking the entire alpha or beta chain, as well as the VJ gene information, and the peptide and MHC data individually. This process not only increased the number of positive samples fivefold but also guided the model to learn the relationships among the various components.
2. Sampling TCRs targeting other pMHCs: we created a 1:1 negative-to-positive sample ratio by randomly disrupting TCR-pMHC pairing. In doing so, we ensured that the distribution of pMHCs in the negative samples mirrored that of the positive samples, thereby preventing the model from learning data biases.
3. Sampling TCRs from public healthy TCR repertoire: We collected a TCR repertoire from PBMCs of healthy individuals (Carter et al., 2019; Grigaityte et al., 2017) and randomly assigned these TCRs to pMHCs, maintaining a 1:1 negative-to-positive ratio.

Through these methods, we aimed to enhance the model’s learning capabilities and improve its predictive performance.

### TcrDesign architecture

#### Transformer encoder

We adopted the standard Transformer encoder architecture, commonly implemented in BERT (Devlin et al., 2018; Vaswani et al., 2017). This neural network architecture employs a self-attention mechanism to process input sequences, effectively capturing the contextual relationships among tokens. This capability allows for parallelization and efficient training, making it particularly suited for generating meaningful, context-aware representations.

We utilize the Transformer encoder to represent the CDR3 sequence of T-cell receptors (TCRs).

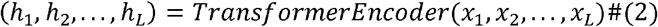

*L* denotes the length of the CDR3 sequence, while (*x*_1_, *x*_2_,..,*x*_*L*_) represents the one-hot encoded amino acids within the CDR3. Additionally, (*h*_1_,*h*_2_,…,*h*_*L*_) ∈ ℝ ^*L* × *hidden_dim*^ are the continuous vectors generated by the Transformer encoder, which represents the latent space of the CDR3 sequence.

#### MLM module

To enable the Transformer encoder to learn the underlying patterns governing TCRs, we employ the MLM training task for pre-training. During this phase, the MLM module learns to reconstruct noisy CDR3 sequences, which are generated by randomly masking 20% of the amino acids in the original sequence. Specifically, among the masked amino acids, 80% are replaced with the special token <mask>, 10% are substituted with random amino acids, and the remaining 10% are left unchanged. This approach enables the model to develop error correction capabilities and learn sequence patterns. The MLM module receives the Transformer encoder output (*h*_1_, *h*_2_, …, *h*_*L*_), and produces a probability distribution over the 20 standard amino acids at the masked positions.

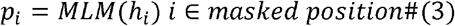

*i* denotes masked positions within the sequence, while *p*_*i*_ ∈ ℝ^20^ represents the probability distribution over the 20 standard amino acids produced by the MLM module. We used cross-entropy as the loss function for MLM. For each masked amino acid in CDR3 sequence, MLM computes its probability distribution and compares it to the true amino acid distribution, which is represented as a one-hot vector.

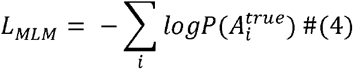

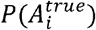 denotes the predicted probability of the actual amino acid.

#### pMHC binding module

For pMHC-BERT, we utilized both an unsupervised masked language modeling (MLM) task and an additional supervised task focused on learning the binding relationship between peptides and MHC during pre-training. The joint training of these two tasks enhances the learned representations of pMHC sequence. This module comprises an mean aggregation module and a multi-layer perceptron (MLP) module, which take representations (*h*_1_, *h*_2_, …,*h*_*L*_) from the output of the Transformer encoder.

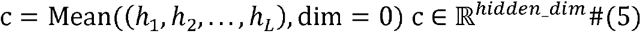

*c* represents the output of the aggregation module, which uses average pooling along the sequence dimension to represent the sequence as a vector of dimension hidden_dim. This vector is subsequently input into the MLP module for processing, resulting in the predicted binding probability value as the output. We used mean squared error (MSE) as the loss function to quantify the difference between the predicted binding probability and the actual binding value.

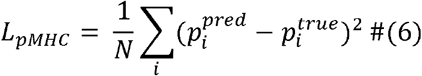

*N* represents the number of training samples, while 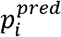 and 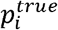 denote the predicted binding probability and the actual binding value, respectively.

#### Details of TcrDesign-B model

We developed the TcrDesign-B model to predict the binding probability between TCRs and pMHC molecules. This model integrates TCR and pMHC information, allowing for missing values. It extracts features using pre-trained BERT models for TCRs and pMHCs, and then integrates those features through a cross-attention layer. The resulting output is processed by a MLP to generate the binding probability. To assess the quality of the outputs from TcrDesign-B, we employed binary cross-entropy (BCE) as the model’s loss function.

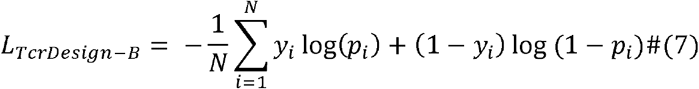

y_i_ is the true label for sample i (0 or 1); p_i_ is the predicted TCR-pMHC binding probability.

#### Details of TcrDesign-G model

We developed the TcrDesign-G model to generate antigen-specific TCRs. The model consists of three modules: the first generates βCDR3 from antigen; the second generates αCDR3 from both the antigen and the βCDR3; and the third generates VJ genes from the generated CDR3 sequences. These modules collaboratively produce a complete TCR. TcrDesign-G employs a Seq2Seq architecture enhanced with an attention mechanism (Bahdanau et al., 2014), which includes Encoder, Attention, and Decoder modules. The encoder initially processes representations (*h*_1_,*h*_2_,…, *h*_*L*_) from pretrained BERT models using a bidirectional GRU layer to obtain hidden states 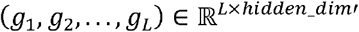.

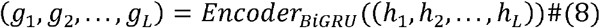

The final hidden state *g*_*L*_ is passed through a linear layer to obtain 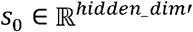, which is then used alongside (*g*_1_, *g*_2_, …, *g*_*L*_) to compute the attention values for each token.

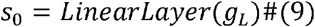

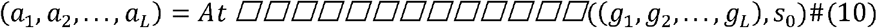

The weighted sum of (*g*_1_, *g*_2_, …, *g*_*L*_) and (*a*_1_,*a*_2_, …, *a*_*L*_) ∈ ℝ^*L*^ produces 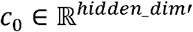, which together with *s*_0_ and the decoder input *x*_1_, guides the generation process of the decoder.

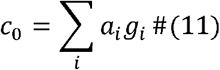

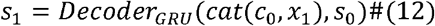

*cat*(*c*_0_,*x*_1_) denotes the concatenation of the two vectors in the feature dimension. This process is repeated based on 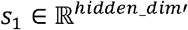, ultimately yielding the decoder output (*s*_1_, *s*_2_, …, *s*_*N*_), where *N* denotes the length of the generated hidden states. After processing through the MLP, a probability distribution over the 20 standard amino acids or the VJ gene vocabulary is produced. The training strategy employs a cross-entropy loss function to predict either the next amino acid or the next VJ gene. During the inference phase, TcrDesign-G utilizes the beam search method to autoregressively generate sequences.

### Training details

We implemented TcrDesign using PyTorch, with a batch size of 512 and training all models on 4 x V100 32GB GPUs. The Transformer encoder is composed of 6 layers and 4 attention heads, featuring an embedding size of 256. We employed grid search to identify optimal mask ratios and learning rates for TCR-BERT and pMHC-BERT. Experimental results demonstrated that a mask ratio of 0.2 and a learning rate of 1×10^-4^ yielded optimal pre-training performance for both models (**Figure S1**). Accordingly, the models were trained for 10 epochs using a learning rate of 1×10^-4^, with a warm-up strategy to stabilize training. For TcrDesign-B, we increased the learning rate to 3×10^-4^ and extended the training to 20 epochs to achieve optimal performance, implementing early stopping with a patience of 3 to prevent overfitting. Similarly, TcrDesign-G was trained for 20 epochs using a learning rate of 3×10^-4^ and a temperature of 1, with the checkpoint achieving the highest average binding score. The GPT baseline (6 layers, 4 attention heads) employed the same training strategy as TcrDesign-G but was trained for 100 epochs due to the absence of pre-training. The input format of the GPT baseline model is <SOS><Epitope><SEP><βCDR3><EOS>. The BERT encoders used for feature extraction in both TcrDesign-B and TcrDesign-G originated from the pretraining phase, with parameters were unfrozen during training.

### Rank score calculation

The binding scores of TCRs to various epitopes, as predicted by TcrDesign-B, may not be directly comparable due to the varying distribution of binding scores for positive TCRs across epitopes. To address this issue and enable cross-epitope comparison of TCR binding strength, we introduced a rank score, inspired from previous studies (Lu et al., 2021; Reynisson et al., 2020). The rank score represents the binding strength between TCRs and pMHCs. To calculate the rank score for a given TCR-pMHC pair, we first established a background distribution by randomly sampling 10,000 TCRs. The binding scores of these TCRs to the same pMHC, as predicted by TcrDesign-B, serve as the background binding distribution. We then determine the percentile rank of the TCR-pMHC pair within this background distribution as rank score. A lower rank score indicates stronger binding, enabling standardized comparison of TCR binding strength across epitopes.

### Sequence similarity

Following the MixTCRpred sequence similarity calculation method (Croce et al., 2024), the sequence similarity between the test epitope and other epitope was determined using the pairwise2 alignment function from the BioPython (Cock et al., 2009) package. The BLOSUM62 scoring matrix (Eddy, 2004) was employed to align the two sequences. The resulting pairwise alignment score is then normalized by dividing it by the score obtained from aligning the test epitope sequence with itself, ensuring that the maximum similarity score is 1. It is important to note that due to the presence of negative values in the BLOSUM62 matrix, the similarity score can also be negative. A score closer to 1 indicates a greater similarity between the peptide pairs.

### Analysis of αTCR and βTCR pairing relationship

Contrastive Language-Image Pretraining (CLIP) is a model developed by OpenAI to learn relationships between text and images. The fundamental concept of the CLIP model is to jointly represent images and text through contrastive learning, aligning them within the same feature space (Radford et al., 2021). In this study, we utilized contrastive learning to train the CLIP model, focusing on aligning the representations of αCDR3 and βCDR3. This was achieved by maximizing the cosine similarity of correct αTCR and βTCR pairs while minimizing the cosine similarity of mismatched pairs, thereby effectively learning the pairing relationships between αTCR and βTCR.

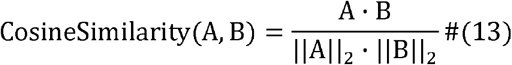

A and B represent feature representations of aCDR3 and bCDR3, respectively, as encoded by the pre-trained BERT model. The term A·B refers to the inner product of the two vectors, while ||A||_2_ and ||B||_2_ indicate the norms (or lengths) of vectors A and B, respectively.

### Epitope-specific TCR clustering

We further investigated the significance of αTCR and βTCR information for the accurate prediction of TcrDesign-B. We selected TCR pairing data for specific antigens and extracted αCDR3 and βCDR3 embeddings after cross-attention in TcrDesign-B. To visualize and cluster this high-dimensional data, we applied UMAP with parameters set to n_neighbors=15 and min_dist=1, reducing the data to two dimensions. We used adjusted rand index (ARI) to quantitatively evaluate clustering performance. KMeans clustering was performed on the data after UMAP dimensionality reduction, and we calculated the ARI to assess the consistency between the clustering results and the actual labels. The ARI ranges from -1 to 1, with higher values indicating greater consistency between the clustering results and the actual labels.

### T-cell line, antigen presenting cells and peptides

HEK-293FT cells were cultured in DMEM (Gibco, cat.C11995500CP) supplemented with 10% FBS (Sigma, cat.F0193) and 100 μg/mL streptomycin and 100 IU/mL penicillin (Gibco, cat. 15140122). Jurkat TCRKO CD8OE cell, a TCR knockout and CD8-stably expressing Jurkat cell clone, is a generous gift from the Chenqi Xu Lab, Shanghai Institute of Biochemistry and Cell Biology, Chinese Academy of Sciences Center for Excellence in Molecular Cell Science. T2 cells (HyCyte, cat.TCH-C419) and Jurkat TCRKO CD8OE cells were grown in RPMI 1640 media supplemented with 10% FBS (Gibco, cat.C11875500CP) and 100 μg/mL streptomycin and 100 IU/mL penicillin (Gibco, cat.15140122). All cells were incubated at 37°C in a 5% CO2 environment. The ELAGIGILTV peptide used in this study was synthesized by GenScript (Nanjing, China). HPLC analysis confirmed peptide purity >98%. ELAGIGILTV peptide was dissolved in DMSO (Solarbio, cat. D8371-100ml) and diluted in RPMI 1640 media supplemented with 10% FBS.

### Lentiviruses production and preparation of TCR-T cells

T cell receptor sequences were constructed based on IMGT reference sequences. V genes, J genes, and CDR3s of the candidate TCRs were synthesized by Shanghai Rui Mian Biological Technology Co.,Ltd. The TCR alpha chain and beta chain were co-expressed on a single plasmid with a P2A cleavage linker. TCR constructs were cloned into pCDH-CMV-MCS-EF1-Puro (Addgene #72265). Lentiviral particles were produced in HEK-293FT cells by co-transfection of packaging vectors pCDH-CMV-MCS-EF1-Puro, psPAX2 (Addgene #12260) and pMD2.G (Addgene #12259) by EZtrans (Shanghai Life iLab Biotech, cat.AC04L099). After 24h, lentivirus particles supernatant was filtered with 0.45 μm syringe filters (Beyotime, cat.FF375-100pcs). For infection, Jurkat TCRKO CD8OE cells were incubated with lentivirus particles supernatant at 37°C overnight and the medium was changed to fresh the next day.

Human peripheral blood mononuclear cells (PBMCs) were purchased from ORIBIOTECH (Shanghai, China). T lymphocytes isolation was performed using EasySep Human CD3+ T Cell Isolation Kit (# 17951, Stem Cell Technologies, Vancouver, Canada) by negative isolation, according to the manufacturer protocols. T lymphocytes were activated using the Human T-activator CD3/CD28 Kit (#11131D, Invitrogen) according to the manufacturer’s instructions. For 1 x 10^6^ T-lymphocytes, 25 μL of Dynabeads coated with anti-CD3 and anti-CD28 antibodies was added to the RPMI 1640 media supplemented with 200U human recombinant IL-2 (#11848-HNAH1-E, Sino Biological). High-titer TCR-lentiviruses were added to transfect primary T cells after activation. After 24 h, the medium was replaced.

### Tetramer staining and flow cytometry

PE-conjugated HLA-A*02:01-ELAGIGILTV recombinant tetramers were obtained from BetterGen (BTG24013100test). Briefly, 10^6^ Jurkat TCRKO CD8OE cells overexpressing candidate TCRs were washed and resuspended in 100 µL of FACS buffer (PBS with 3% FBS). Subsequently, 1 µL of tetramer was added to the cells, which were then incubated in the dark on ice for 1h. This was followed by staining with 1 µL of APC anti-human TCR α/β antibody (BioLegend, cat.306717) in the dark on ice for an additional 30 minutes to detect surface TCRs. Samples were washed twice with FACS buffer and then resuspended in 200 µL of fixative solution (PBS + 1% paraformaldehyde) prior to analysis using a BD LSRFortessa X20 flow cytometer.

### T-cell activation assay

To assess the activation of a TCR against an antigen of interest, we developed a Jurkat-NFAT-ZsGreen reporter system for the nuclear factor of activated T cells (NFAT). For every 10^6^ candidate TCR-overexpressing CD8+ Jurkat cells (described above) were electroporated with 2 μg of pEF1a-NFAT-ZsGreen reporter plasmid using the Lonza 4D-Nucleofector. The Jurkat-NFAT-ZsGreen reporter cells were co-cultured with T2 antigen-presenting cells expressing HLA-A*02:01 at 37°C, 5% CO2. The T2 cells were pulsed for 2h with 10 μg/mL ELAGIGILTV peptide, then incubated at a 1:1 ratio with Jurkat-NFAT-ZsGreen reporter cells. After 24h, CD69 and GFP levels of Jurkat cells were analyzed by BD LSRFortessa X20 flow cytometry. CD69 levels were measured with PE anti-human CD69 antibody (Biolegend, cat.310905), TCR expression levels were measured with APC anti-human TCR α/β antibody (BioLegend, cat.306717). Flow cytometry analysis was performed using FlowJo 10.8 **(Figure S18)**.

### IL-2 ELISA assay

To measure the IL-2 level in candidate TCR-transduced Jurkat T cells that were activated by antigen presenting cell, 1 × 10^5^ Jurkat cells were seeded in 96-well plates and treated with 1 × 10^5^ serially diluted peptides pulsed T2 cells for 24 h. Cells were centrifuged at 500 × g for 5 min at RT, and the supernatants were collected. IL-2 was quantitated with ELISA Kit (MA0726, MeilunBio). The assays were carried out in accordance with the manufacturer’s instructions.

### Bioluminescence-based cytotoxicity assay

Target cell T2-luciferase were generated via lentiviral transduction. 25,000 target cells in T cell medium were seeded in 96-well plates co-cultured with ELApositive TCR-T, ELA3 TCR-T, ELA4 TCR-T, ELA6 TCR-T and mock-T cells at varying effector-to-target (E:T) ratios for 24 h in triplicate. Wells containing target cells with mock-T cells served as negative controls for baseline cytotoxicity. Subsequently, the supernatant was removed and the collected cells were lysed, centrifuged at 800×g for 5□min to obtain the supernatant. The luciferase assay kit (RG005, Beyotime Biotechnology Co., Ltd.) was used to measure luciferase activity according to the instructions.

The percent (%) specific lysis of tumor cells was calculated using the formula: % specific lysis = (luminescence of tumor cells cocultured with mock - T cells luminescence of tumor cells co-cultured with TCR-T cells) / luminescence of tumor cell line cocultured with mock-T cells × 100.

## Statistical analyses

All significance tests were two-sided, with a P value of less than 0.05 considered statistically significant (ns P > 0.05, *P < 0.05, **P < 0.01, ***P < 0.001). Performance metrics, including AUC, PPV, and ARI, along with KMeans clustering, were computed using the Python package scikit-learn v1.10.1. UMAP was implemented using the Python package umap-learn v0.5.4. Sequence similarity was calculated with Biopython v1.83, while sequence motifs were visualized using the Python package logomaker v0.8, which was configured with custom color schemes. The open-source software PyMOL was utilized for visualizing the 3D structure of TCR-pMHC complexes.

## Data and code availability

All data is sourced from public databases such as VDJdb, McPAS, IEDB, TCR3d, and TCRdb. The data and weights used by TcrDesign can be downloaded from zenodo website at https://zenodo.org/records/14545852. The TcrDesign code repository, which includes detailed installation and usage instructions, is available on https://github.com/XSLiuLab/TcrDesign.

## Acknowledgement

We thank ShanghaiTech University High Performance Computing Public Service Platform for computing services. We thank multi-omics facility, molecular and cell biology core facility of ShanghaiTech University for technical help. This work is supported by Shanghai Science and Technology Commission (24J22800700), cross disciplinary Research Fund of Shanghai Ninth People’s Hospital, Shanghai JiaoTong University School of Medicine (JYJC202227), National Natural Science Foundation of China (82373149), open project fund of the National Health Commission’s key laboratory of individualized diagnosis and treatment of nasopharyngeal cancer (2023NPCCK02), Shanghai Municipal Health Commission (2024CXJQ02), and startup funding from ShanghaiTech University.

## Author contributions

X.L. designed the study. K.D., J.C. X.L. wrote the manuscript. K.D., J.C. developed the TcrDesign tool. J.C. performed the experiments validating the function of AI generated TCRs. X.Z. participated in experimental validations. K.D., J.C., D.Q., T.W. participated in the computational analysis. H.W. provided experimental materials and participated in critical discussions.

## References

Bagaev, D.V., Vroomans, R.M.A., Samir, J., Stervbo, U., Rius, C., Dolton, G., Greenshields-Watson, A., Attaf, M., Egorov, E.S., Zvyagin, I.V., et al. (2020). VDJdb in 2019: database extension, new analysis infrastructure and a T-cell receptor motif compendium. Nucleic Acids Research 48, D1057–D1062.

Bahdanau, D., Cho, K., and Bengio, Y. (2014). Neural Machine Translation by Jointly Learning to Align and Translate. arXiv 1409.0473.

Baulu, E., Gardet, C., Chuvin, N., and Depil, S. (2023). TCR-engineered T cell therapy in solid tumors: State of the art and perspectives. Science Advances 9, eadf3700.

Benayas, A., Sicilia, M.A., and Mora-Cantallops, M. (2024). A comparative analysis of encoder only and decoder only models in intent classification and sentiment analysis: Navigating the trade-offs in model size and performance. In Review 10.21203/rs.3.rs-3865391/v1.

Bennett, N.R., Watson, J.L., Ragotte, R.J., Borst, A.J., See, D.L., Weidle, C., Biswas, R., Yu, Y., Shrock, E.L., Ault, R., et al. (2025). Atomically accurate de novo design of antibodies with RFdiffusion. bioRxiv 585103.

Bentzen, A.K., Marquard, A.M., Lyngaa, R., Saini, S.K., Ramskov, S., Donia, M., Such, L., Furness, A.J.S., McGranahan, N., Rosenthal, R., et al. (2016). Large-scale detection of antigen-specific T cells using peptide-MHC-I multimers labeled with DNA barcodes. Nature Biotechnology 34, 1037–1045.

Brady, B.L., Steinel, N.C., and Bassing, C.H. (2010). Antigen Receptor Allelic Exclusion: An Update and Reappraisal. The Journal of Immunology 185, 3801–3808.

Brightman, S.E., Naradikian, M.S., Miller, A.M., and Schoenberger, S.P. (2020). Harnessing neoantigen specific CD4 T cells for cancer immunotherapy. Journal of Leukocyte Biology 107, 625–633.

Carter, J.A., Preall, J.B., Grigaityte, K., Goldfless, S.J., Jeffery, E., Briggs, A.W., Vigneault, F., and Atwal, G.S. (2019). Single T Cell Sequencing Demonstrates the Functional Role of αβ TCR Pairing in Cell Lineage and Antigen Specificity. Frontiers in Immunology 10, 1516.

Chen, S.-Y., Yue, T., Lei, Q., and Guo, A.-Y. (2021). TCRdb: a comprehensive database for T-cell receptor sequences with powerful search function. Nucleic Acids Research 49, D468–D474.

Cock, P.J.A., Antao, T., Chang, J.T., Chapman, B.A., Cox, C.J., Dalke, A., Friedberg, I., Hamelryck, T., Kauff, F., Wilczynski, B., et al. (2009). Biopython: freely available Python tools for computational molecular biology and bioinformatics. Bioinformatics 25, 1422–1423.

Croce, G., Bobisse, S., Moreno, D.L., Schmidt, J., Guillame, P., Harari, A., and Gfeller, D. (2024). Deep learning predictions of TCR-epitope interactions reveal epitope-specific chains in dual alpha T cells. Nature Communications 15, 3211.

Dash, P., Fiore-Gartland, A.J., Hertz, T., Wang, G.C., Sharma, S., Souquette, A., Crawford, J.C., Clemens, E.B., Nguyen, T.H.O., Kedzierska, K., et al. (2017). Quantifiable predictive features define epitope-specific T cell receptor repertoires. Nature 547, 89–93.

Davis, M.M., and Bjorkman, P.J. (1988). T-cell antigen receptor genes and T-cell recognition. Nature 334, 395–402.

Deng, L., Ly, C., Abdollahi, S., Zhao, Y., Prinz, I., and Bonn, S. (2023). Performance comparison of TCR-pMHC prediction tools reveals a strong data dependency. Frontiers in Immunology 14, 1128326.

Devlin, J., Chang, M.-W., Lee, K., and Toutanova, K. (2018). BERT: Pre-training of Deep Bidirectional Transformers for Language Understanding. arXiv 1810.04805.

Dines, J.N., Manley, T.J., Svejnoha, E., Simmons, H.M., Taniguchi, R., Klinger, M., Baldo, L., and Robins, H. (2020). The ImmuneRACE Study: A Prospective Multicohort Study of Immune Response Action to COVID-19 Events with the ImmuneCODE™ Open Access Database. 10.1101/2020.08.17.20175158.

Dolton, G., Tungatt, K., Lloyd, A., Bianchi, V., Theaker, S.M., Trimby, A., Holland, C.J., Donia, M., Godkin, A.J., Cole, D.K., et al. (2015). More tricks with tetramers: a practical guide to staining T cells with peptide– MHC multimers. Immunology 146, 11–22.

Dossa, R.G., Cunningham, T., Sommermeyer, D., Medina-Rodriguez, I., Biernacki, M.A., Foster, K., and Bleakley, M. (2018). Development of T-cell immunotherapy for hematopoietic stem cell transplantation recipients at risk of leukemia relapse. Blood 131, 108–120.

Eddy, S.R. (2004). Where did the BLOSUM62 alignment score matrix come from? Nature Biotechnology 22, 1035–1036.

Fast, E., Dhar, M., and Chen, B. (2023). TAPIR: a T-cell receptor language model for predicting rare and novel targets. bioRxiv 557285.

Gfeller, D., Schmidt, J., Croce, G., Guillaume, P., Bobisse, S., Genolet, R., Queiroz, L., Cesbron, J., Racle, J., and Harari, A. (2023). Improved predictions of antigen presentation and TCR recognition with MixMHCpred2.2 and PRIME2.0 reveal potent SARS-CoV-2 CD8+ T-cell epitopes. Cell Systems 14, 72–83.e75.

Grazioli, F., Mösch, A., Machart, P., Li, K., Alqassem, I., O’Donnell, T.J., and Min, M.R. (2022). On TCR binding predictors failing to generalize to unseen peptides. Frontiers in Immunology 13, 1014256.

Grigaityte, K., Carter, J.A., Goldfless, S.J., Jeffery, E.W., Hause, R.J., Jiang, Y., Koppstein, D., Briggs, A.W., Church, G.M., Vigneault, F., et al. (2017). Single-cell sequencing reveals αβ chain pairing shapes the T cell repertoire. bioRxiv 213462.

Han, A., Glanville, J., Hansmann, L., and Davis, M.M. (2014). Linking T-cell receptor sequence to functional phenotype at the single-cell level. Nature Biotechnology 32, 684–692.

Harris, Daniel T., Singh, Nishant K., Cai, Q., Smith, Sheena N., Vander Kooi, Craig W., Procko, E., Kranz, David M., and Baker, Brian M. (2016). An Engineered Switch in T Cell Receptor Specificity Leads to an Unusual but Functional Binding Geometry. Structure 24, 1142–1154.

Heath, W.R., Carbone, F.R., Bertolino, P., Kelly, J., Cose, S., and Miller, J.F.A.P. (1995). Expression of two T cell receptor α chains on the surface of normal murine T cells. European Journal of Immunology 25, 1617–1623.

Hudson, D., Fernandes, R.A., Basham, M., Ogg, G., and Koohy, H. (2023). Can we predict T cell specificity with digital biology and machine learning? Nature Reviews Immunology 23, 511–521.

Isacchini, G., Walczak, A.M., Mora, T., and Nourmohammad, A. (2021). Deep generative selection models of T and B cell receptor repertoires with soNNia. Proceedings of the National Academy of Sciences 118, e2023141118.

Jensen, M.F., and Nielsen, M. (2023). NetTCR 2.2 - Improved TCR specificity predictions by combining pan- and peptide-specific training strategies, loss-scaling and integration of sequence similarity. bioRxiv 562001.

Joglekar, A.V., and Li, G. (2021). T cell antigen discovery. Nature Methods 18, 873–880.

Krogsgaard, M., and Davis, M.M. (2005). How T cells ‘see’ antigen. Nature Immunology 6, 239–245.

Leclercq, G., Steinhoff, N., Haegel, H., De Marco, D., Bacac, M., and Klein, C. (2022). Novel strategies for the mitigation of cytokine release syndrome induced by T cell engaging therapies with a focus on the use of kinase inhibitors. Oncoimmunology 11, 2083479.

Legut, M., Dolton, G., Mian, A.A., Ottmann, O.G., and Sewell, A.K. (2018). CRISPR-mediated TCR replacement generates superior anticancer transgenic T cells. Blood 131, 311–322.

Lin, Y., Zhang, D., and Liu, Y. (2024). TCR-GPT: Integrating Autoregressive Model and Reinforcement Learning for T-Cell Receptor Repertoires Generation. arXiv 2408.01156.

Lu, T., Zhang, Z., Zhu, J., Wang, Y., Jiang, P., Xiao, X., Bernatchez, C., Heymach, J.V., Gibbons, D.L., Wang, J., et al. (2021). Deep learning-based prediction of the T cell receptor–antigen binding specificity. Nature Machine Intelligence 3, 864–875.

Malu, S., Malshetty, V., Francis, D., and Cortes, P. (2012). Role of non-homologous end joining in V(D)J recombination. Immunologic Research 54, 233–246.

Mayer-Blackwell, K., Schattgen, S., Cohen-Lavi, L., Crawford, J.C., Souquette, A., Gaevert, J.A., Hertz, T., Thomas, P.G., Bradley, P., and Fiore-Gartland, A. (2021). TCR meta-clonotypes for biomarker discovery with tcrdist3 enabled identification of public, HLA-restricted clusters of SARS-CoV-2 TCRs. eLife 10, e68605.

Meysman, P., Barton, J., Bravi, B., Cohen-Lavi, L., Karnaukhov, V., Lilleskov, E., Montemurro, A., Nielsen, M., Mora, T., Pereira, P., et al. (2023). Benchmarking solutions to the T-cell receptor epitope prediction problem: IMMREP22 workshop report. ImmunoInformatics 9, 100024.

Montemurro, A., Jessen, L.E., and Nielsen, M. (2022). NetTCR-2.1: Lessons and guidance on how to develop models for TCR specificity predictions. Frontiers in Immunology 13, 1055151.

Moravec, Z., Zhao, Y., Voogd, R., Cook, D.R., Kinrot, S., Capra, B., Yang, H., Raud, B., Ou, J., Xuan, J., et al. (2024). Discovery of tumor-reactive T cell receptors by massively parallel library synthesis and screening. Nature Biotechnology 43, 214–222.

Moris, P., De Pauw, J., Postovskaya, A., Gielis, S., De Neuter, N., Bittremieux, W., Ogunjimi, B., Laukens, K., and Meysman, P. (2021). Current challenges for unseen-epitope TCR interaction prediction and a new perspective derived from image classification. Briefings in Bioinformatics 22, bbaa318.

Nielsen, M., Eugster, A., Jensen, M.F., Goel, M., Tiffeau-Mayer, A., Pelissier, A., Valkiers, S., Martínez, M.R., Meynard-Piganeeau, B., Greiff, V., et al. (2024). Lessons learned from the IMMREP23 TCR-epitope prediction challenge. ImmunoInformatics 16, 100045.

Nilsson, J.B., Kaabinejadian, S., Yari, H., Kester, M.G.D., Van Balen, P., Hildebrand, W.H., and Nielsen, M. (2023). Accurate prediction of HLA class II antigen presentation across all loci using tailored data acquisition and refined machine learning. Science Advances 9, eadj6367.

Nolan, S., Vignali, M., Klinger, M., Dines, J.N., Kaplan, I.M., Svejnoha, E., Craft, T., Boland, K., Pesesky, M., Gittelman, R.M., et al. (2020). A large-scale database of T-cell receptor beta (TCRβ) sequences and binding associations from natural and synthetic exposure to SARS-CoV-2. In Review 10.21203/rs.3.rs-51964/v1.

O’Donnell, T.J., Rubinsteyn, A., and Laserson, U. (2020). MHCflurry 2.0: Improved Pan-Allele Prediction of MHC Class I-Presented Peptides by Incorporating Antigen Processing. Cell Systems 11, 42–48.e47.

Padovan, E., Casorati, G., Dellabona, P., Meyer, S., Brockhaus, M., and Lanzavecchia, A. (1993). Expression of Two T Cell Receptor α Chains: Dual Receptor T Cells. Science 262, 422–424.

Radford, A., Kim, J.W., Hallacy, C., Ramesh, A., Goh, G., Agarwal, S., Sastry, G., Askell, A., Mishkin, P., Clark, J., et al. (2021). Learning Transferable Visual Models From Natural Language Supervision. arXiv 2103.00020.

Raffel, C., Shazeer, N., Roberts, A., Lee, K., Narang, S., Matena, M., Zhou, Y., Li, W., and Liu, P.J. (2023). Exploring the limits of transfer learning with a unified text-to-text transformer. arXiv 1910.10683.

Reynisson, B., Alvarez, B., Paul, S., Peters, B., and Nielsen, M. (2020). NetMHCpan-4.1 and NetMHCIIpan-4.0: improved predictions of MHC antigen presentation by concurrent motif deconvolution and integration of MS MHC eluted ligand data. Nucleic acids research 48, W449–W454.

Rodrigues Dos Santos, A., Zanini, D., and Andolfatto, D. (2024). Cytokine release syndrome after chimeric antigen receptor T cell therapy in patients with diffuse large B-cell lymphoma: a systematic review. Hematol Transfus Cell Ther 46 Suppl 6, S306–S315.

Schatz, D.G., and Ji, Y. (2011). Recombination centres and the orchestration of V(D)J recombination. Nature Reviews Immunology 11, 251–263.

Schuldt, N.J., and Binstadt, B.A. (2019). Dual TCR T Cells: Identity Crisis or Multitaskers? The Journal of Immunology 202, 637–644.

Shao, Y.C., Mu, Q.D., Wang, R., Luo, H.B., Song, Z.J., Wang, P.F., Song, J.S., Ge, C.D., Zhang, J.Y., Min, J.X., et al. (2025). SLC39A10 is a key zinc transporter in T cells and its loss mitigates autoimmune disease. Science China-Life Sciences 68, 1855–1870.

Shi, Y., Parks, J.M., and Smith, J.C. (2025). Comparative Analysis of TCR and TCR-pMHC Complex Structure Prediction Tools. Journal of chemical information and modeling 65, 7156–7173.

Shugay, M., Bagaev, D.V., Zvyagin, I.V., Vroomans, R.M., Crawford, J.C., Dolton, G., Komech, E.A., Sycheva, A.L., Koneva, A.E., Egorov, E.S., et al. (2018). VDJdb: a curated database of T-cell receptor sequences with known antigen specificity. Nucleic Acids Research 46, D419–D427.

Spindler, M.J., Nelson, A.L., Wagner, E.K., Oppermans, N., Bridgeman, J.S., Heather, J.M., Adler, A.S., Asensio, M.A., Edgar, R.C., Lim, Y.W., et al. (2020). Massively parallel interrogation and mining of natively paired human TCRαβ repertoires. Nature Biotechnology 38, 609–619.

Springer, I., Besser, H., Tickotsky-Moskovitz, N., Dvorkin, S., and Louzoun, Y. (2020). Prediction of Specific TCR-Peptide Binding From Large Dictionaries of TCR-Peptide Pairs. Frontiers in Immunology 11, 1803.

Springer, I., Tickotsky, N., and Louzoun, Y. (2021). Contribution of T Cell Receptor Alpha and Beta CDR3, MHC Typing, V and J Genes to Peptide Binding Prediction. Frontiers in Immunology 12, 664514.

Stone, J.D., and Kranz, D. (2013). Role of T cell receptor affinity in the efficacy and specificity of adoptive T cell therapies. Frontiers in Immunology 4, 244.

Su, D., and Liu, G. (2025). Targeted Ablation of Canonical Disulfide Bonds Mitigates TCR Mismatching in Engineered T Cells. bioRxiv 656745.

Su, J., Han, C., Zhou, Y., Shan, J., Zhou, X., and Yuan, F. (2023). SaProt: Protein Language Modeling with Structure-aware Vocabulary. bioRxiv 560349.

Sun, J., Zhu, T., Cui, Y., and Wu, B. (2025). Structure-based self-supervised learning enables ultrafast protein stability prediction upon mutation. The Innovation 6, 100750.

Sussman, J.L., Lin, D., Jiang, J., Manning, N.O., Prilusky, J., Ritter, O., and Abola, E.E. (1998). Protein Data Bank (PDB): Database of Three-Dimensional Structural Information of Biological Macromolecules. Acta Crystallographica Section D Biological Crystallography 54, 1078–1084.

Tanno, H., Gould, T.M., McDaniel, J.R., Cao, W., Tanno, Y., Durrett, R.E., Park, D., Cate, S.J., Hildebrand, W.H., Dekker, C.L., et al. (2020). Determinants governing T cell receptor α/β-chain pairing in repertoire formation of identical twins. Proceedings of the National Academy of Sciences 117, 532–540.

Tickotsky, N., Sagiv, T., Prilusky, J., Shifrut, E., and Friedman, N. (2017). McPAS-TCR: a manually curated catalogue of pathology-associated T cell receptor sequences. Bioinformatics 33, 2924–2929.

Van Gent, D.C., Ramsden, D.A., and Gellert, M. (1996). The RAG1 and RAG2 Proteins Establish the 12/23 Rule in V(D)J Recombination. Cell 85, 107–113.

Vaswani, A., Shazeer, N., Parmar, N., Uszkoreit, J., Jones, L., Gomez, A.N., Kaiser, L., and Polosukhin, I. (2017). Attention Is All You Need. arXiv 1706.03762.

Vita, R., Mahajan, S., Overton, J.A., Dhanda, S.K., Martini, S., Cantrell, J.R., Wheeler, D.K., Sette, A., and Peters, B. (2019). The Immune Epitope Database (IEDB): 2018 update. Nucleic Acids Research 47, D339–D343.

Wang, X., Song, X., Li, Y., Ding, Y., Yin, C., Ren, T., and Zhang, W. (2025). Integrated system for screening tumor-specific TCRs, epitopes, and HLA subtypes using single-cell sequencing data. J Immunother Cancer 13, e012029.

Xiong, C., Huang, L., Kou, H., Wang, C., Zeng, X., Sun, H., Liu, S., Wu, B., Li, J., Wang, X., et al. (2022). Identification of novel HLA-A*11:01-restricted HPV16 E6/E7 epitopes and T-cell receptors for HPV-related cancer immunotherapy. J Immunother Cancer 10, e004790.

Xu, H., Li, L., Pan, S., Cheng, P., Wang, Y., Rong, Z., Liu, F., Huang, X., Wang, S., and Shu, W. (2026). Protein foundation models: a comprehensive survey. SCIENCE CHINA Life Sciences 10.1007/s11427-025-3147-2.

Xu, R., Du, S., Zhu, J., Meng, F., and Liu, B. (2022). Neoantigen-targeted TCR-T cell therapy for solid tumors: How far from clinical application. Cancer Letters 546, 215840.

Yang, J., He, B., Zhao, Y., Jiang, F., Wang, Z., Guo, Y., Xu, Z., Yuan, B., Song, J., Zhang, Q., et al. (2023). De novo generation of T-cell receptors with desired epitope-binding property by leveraging a pre-trained large language model. bioRxiv 562845.

Yin, R., Ribeiro-Filho, H.V., Lin, V., Gowthaman, R., Cheung, M., and Pierce, Brian G. (2023). TCRmodel2: high-resolution modeling of T cell receptor recognition using deep learning. Nucleic Acids Research 51, W569–W576.

Zhang, L., Pang, H., Zhang, C., Li, S., Tan, Y., Jiang, F., Li, M., Yu, Y., Zhou, Z., Wu, B., et al. (2025). VenusMutHub: A systematic evaluation of protein mutation effect predictors on small-scale experimental data. Acta Pharmaceutica Sinica B 6, S2211383525001650.

Zhang, W., Hawkins, P.G., He, J., Gupta, N.T., Liu, J., Choonoo, G., Jeong, S.W., Chen, C.R., Dhanik, A., Dillon, M., et al. (2021). A framework for highly multiplexed dextramer mapping and prediction of T cell receptor sequences to antigen specificity. Science Advances 7, eabf5835.

Zhang, W., Wang, L., Liu, K., Wei, X., Yang, K., Du, W., Wang, S., Guo, N., Ma, C., Luo, L., et al. (2019). PIRD: Pan Immune Repertoire Database. Bioinformatics 36, 897–903.

Zhong, S., Malecek, K., Johnson, L.A., Yu, Z., Vega-Saenz de Miera, E., Darvishian, F., McGary, K., Huang, K., Boyer, J., Corse, E., et al. (2013). T-cell receptor affinity and avidity defines antitumor response and autoimmunity in T-cell immunotherapy. Proc Natl Acad Sci U S A 110, 6973–6978.

Zhou, W., Xiang, W., Yu, J., Ruan, Z., Pan, Y., Wang, K., and Liu, J. (2024a). NeoTCR: an immunoinformatic database of experimentally-supported functional neoantigen-specific TCR sequences. Genomics, Proteomics & Bioinformatics 23, qzae010.

Zhou, Z., Chen, J., Lin, S., Hong, L., Wei, D.-Q., and Xiong, Y. (2024b). GRATCR: epitope-specific T cell receptor sequence generation with data-efficient pre-trained models. bioRxiv 604503.

