## Supplemental File for "TcrDesign: De novo design of epitope specific full-length T cell receptors"

***Supplementary Materials***

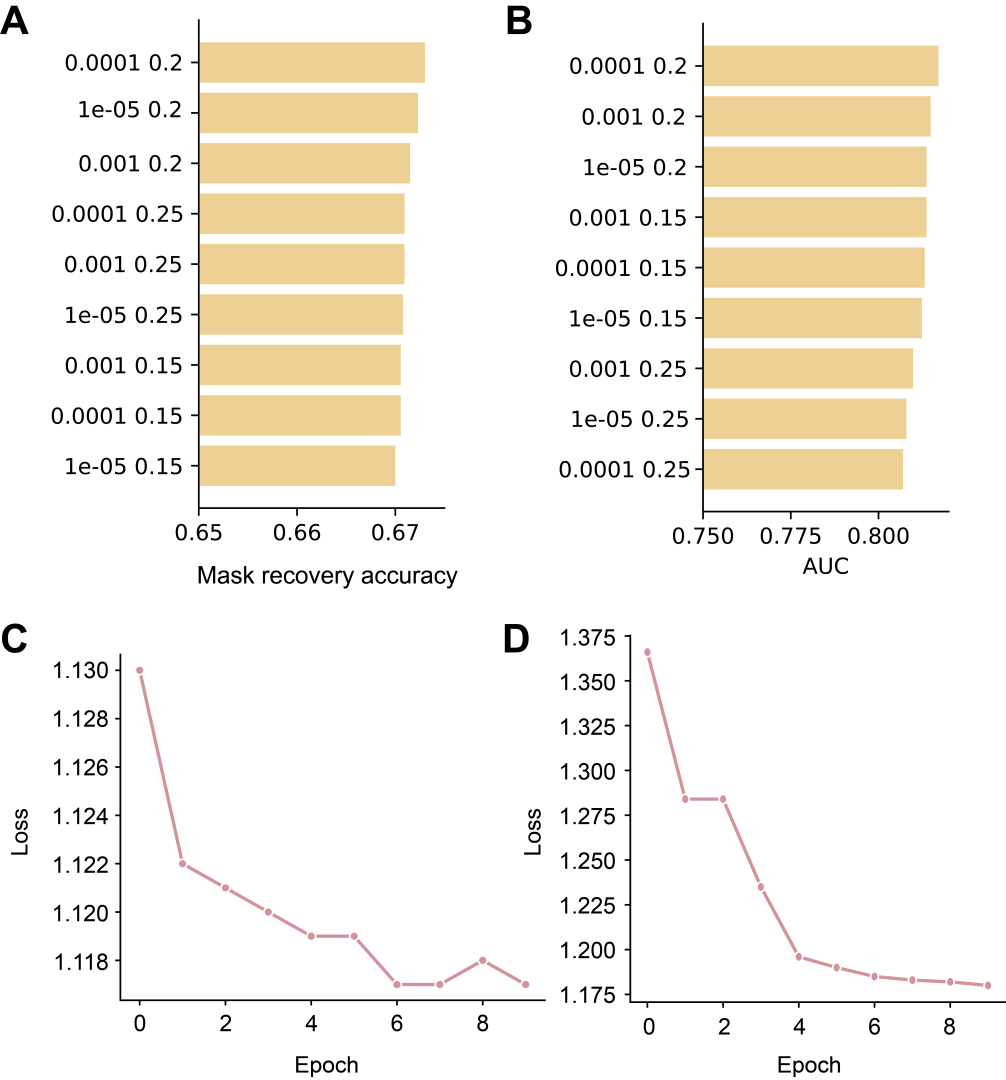

**Figure S1. Pre-training details.** (**A**) Hyperparameter search for bCDR3-BERT, evaluated using masked token recovery rate. (**B**) Hyperparameter search for pMHC-BERT, evaluated using the AUC of peptide-MHC binding prediction. (**C**) Training curves of bCDR3-BERT. (**D**) Training curves of pMHC-BERT.

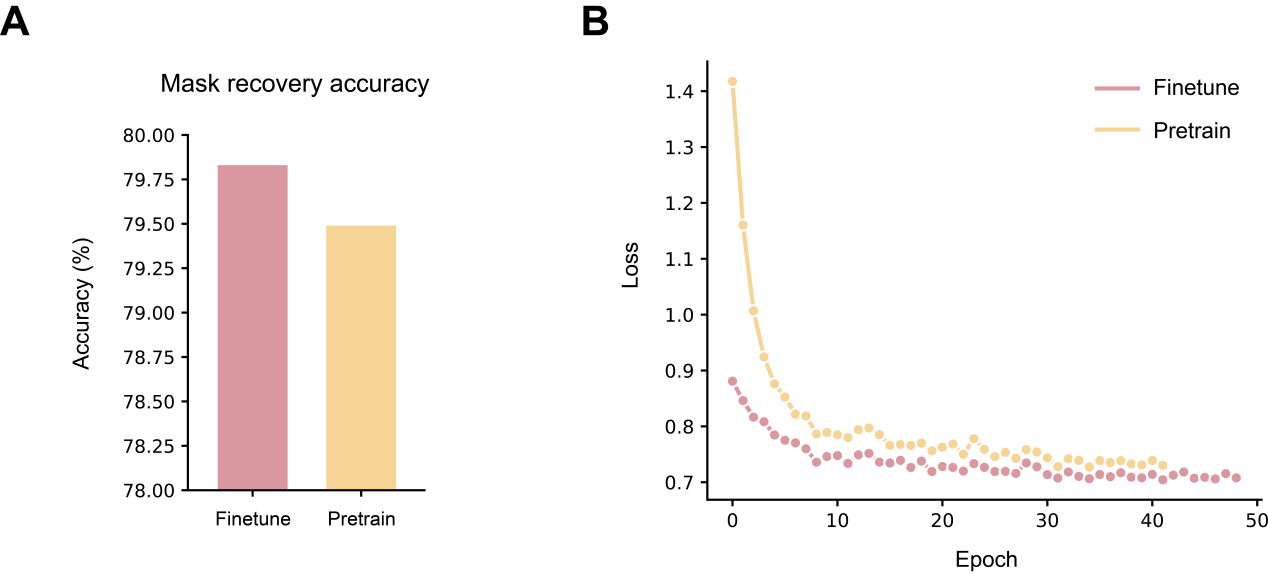

**Figure S2. Training evaluation of αCDR3-BERT.** (**A**) Comparison of masked recovery accuracy on the αCDR3 validation set for fine-tuning based on βCDR3-BERT and pre-training (retraining). (**B**) Training process of fine-tuning and pre-training models.

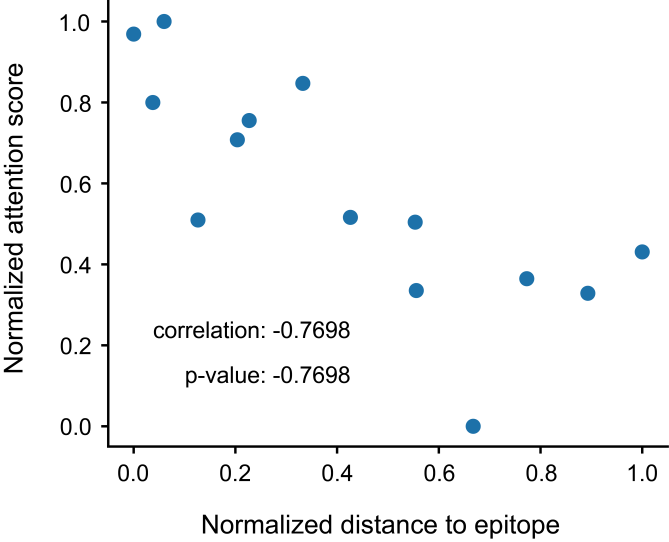

**Figure S3. Correlation analysis for one pair of βCDR3-epitope.** To explore the biological significance of attention weights in TCR-BERT, we analyzed a pair of βCDR3-epitope as an example. βCDR3-BERT extracts the attention weights for each amino acid in the βCDR3 sequence and computes the distance from each amino acid in βCDR3 to the epitope based on structural calculations, followed by a correlation analysis.

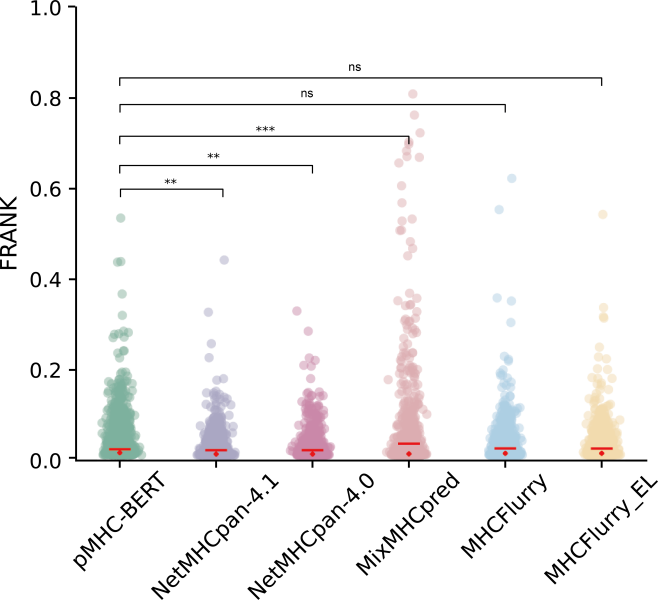

**Figure S4. Comparison of the prediction performance of pMHC-BERT against other models on the CD8 epitope dataset.**

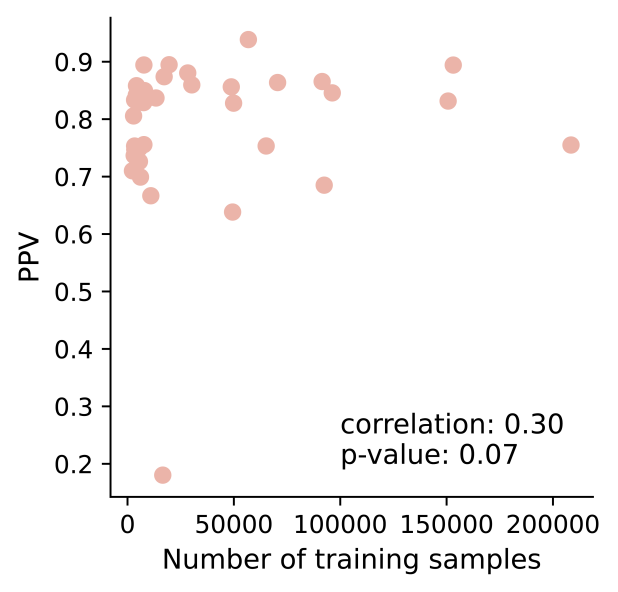

**Figure S5. The scatter between the number of training samples and PPV across different MHC subtypes, each dot represent a MHC subtype. Spearman correlation was calculated.**

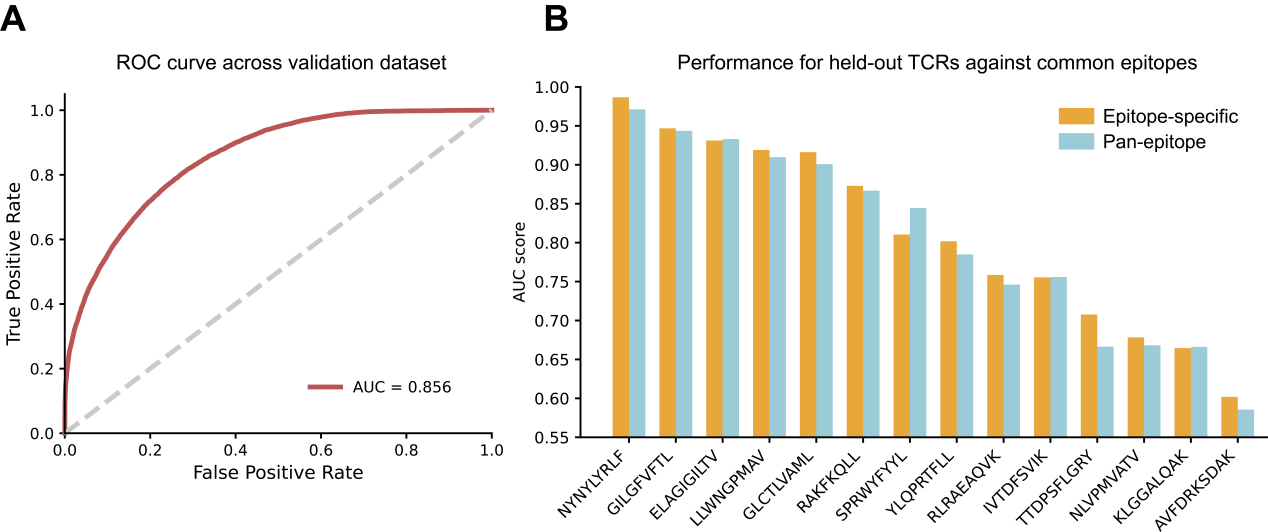

**Figure S6. Performance of two versions (epitope-specific and pan-epitope) of TcrDesign-B.** (**A**) The AUC curve of pan-epitope TcrDesign-B on the validation dataset. (**B**) The AUC scores of two versions of TcrDesign-B on the independent common epitope dataset.

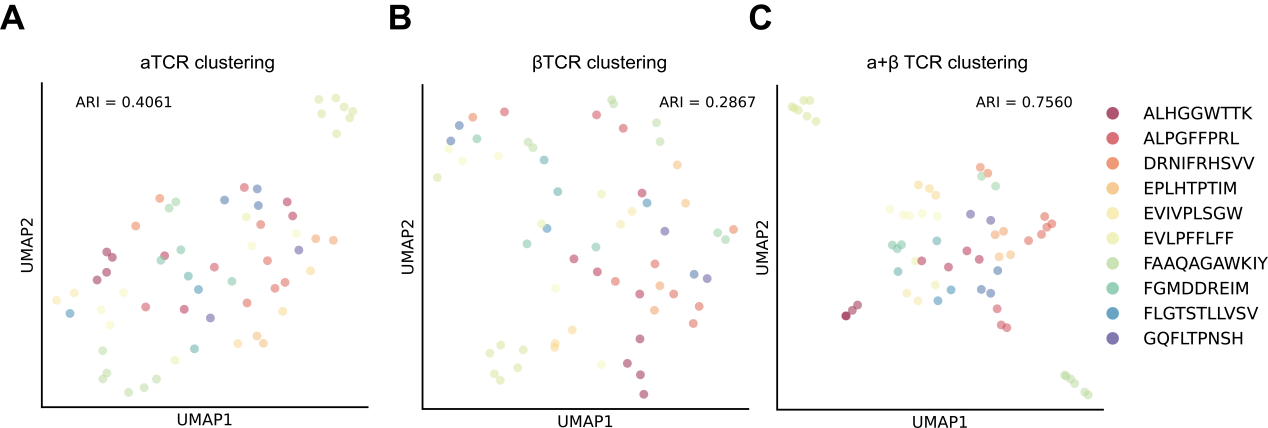

**Figure S7. Visualization of TCRs on the novel epitope dataset.** UMAP dimensionality reduction of epitope-specific TCRs from the novel epitope dataset utilizing TCR embeddings extracted by TcrDesign-B. The ARI was employed to evaluate clustering performance, where higher values indicate better clustering.

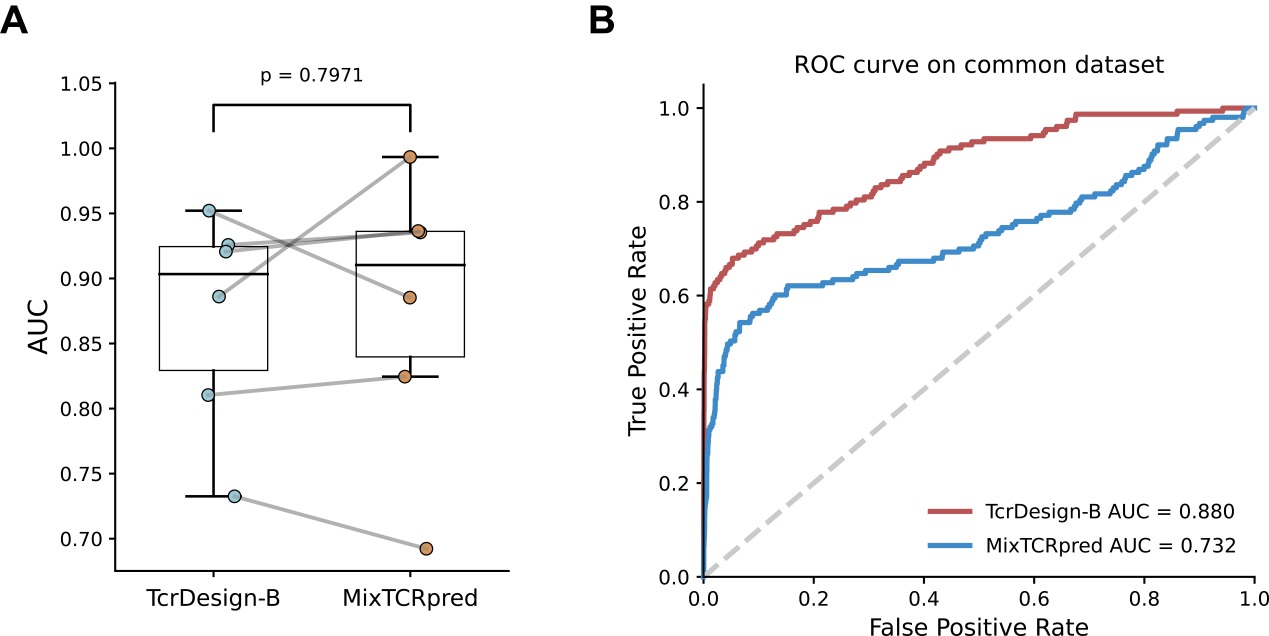

**Figure S8. Performance of TcrDesign-B and MixTCRpred.** (**A**) AUC scores of TcrDesign-B and MixTCRpred for each common epitope. (**B**) ROC curve of TcrDesign-B and MixTCRpred on the common dataset.

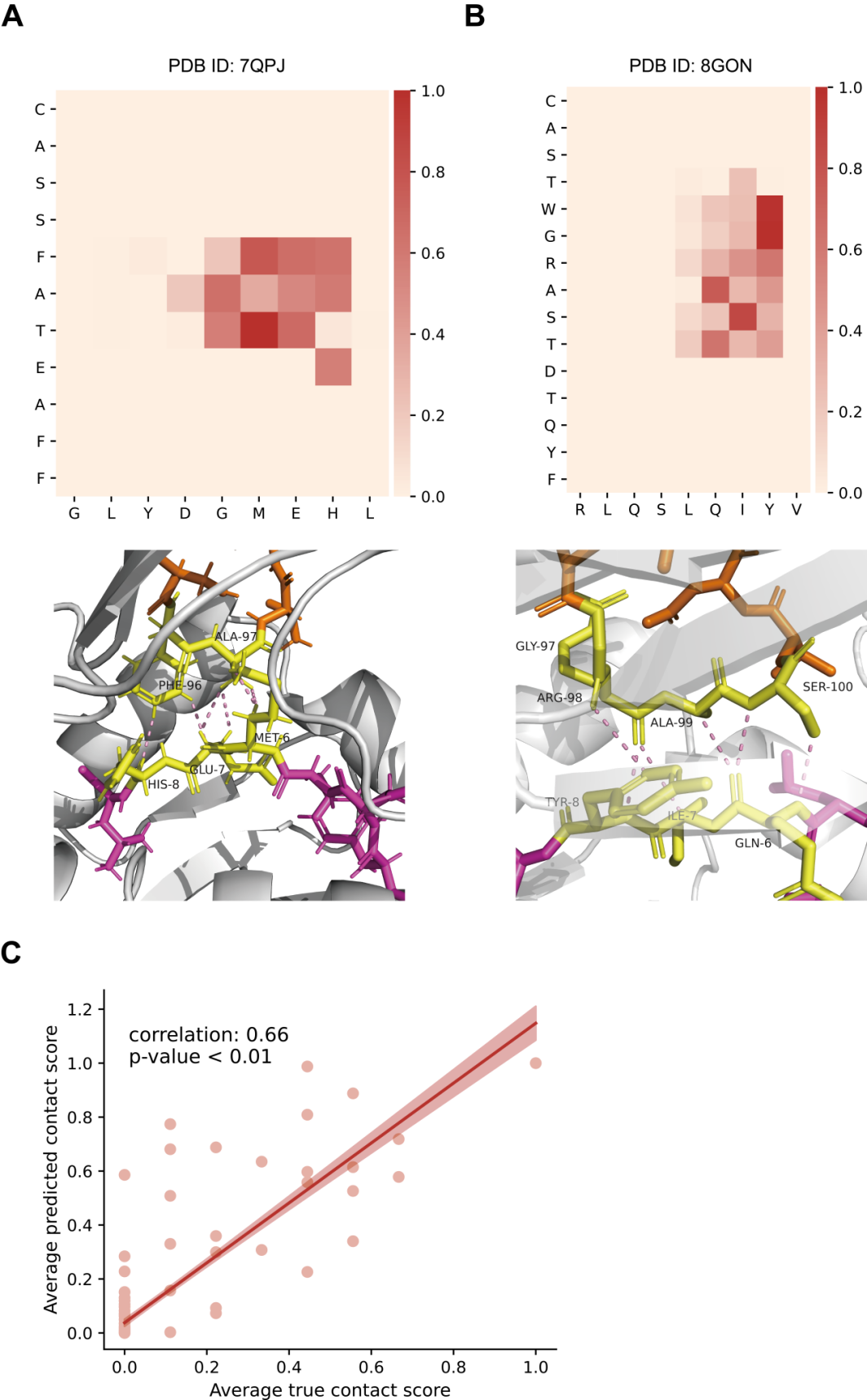

**Figure S9. TcrDesign-B predicts TCR-pMHC interactions at amino acid level.** Extended TcrDesign-B predicts the epitope-βCDR3 interaction maps for two examples (**A**: 7QPJ, **B** : 8GON). (**C**) The Pearson correlation coefficient calculated between the predicted contact scores and the actual contact scores (normalized Ca-Ca distances).

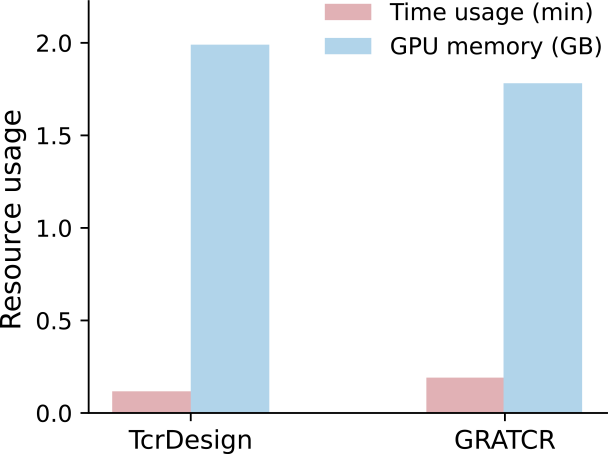

**Figure S10. Time consumption and GPU memory usage for generating 1,000 TCRs targeting ELAGIGILTV by TcrDesign and GRATCR.**

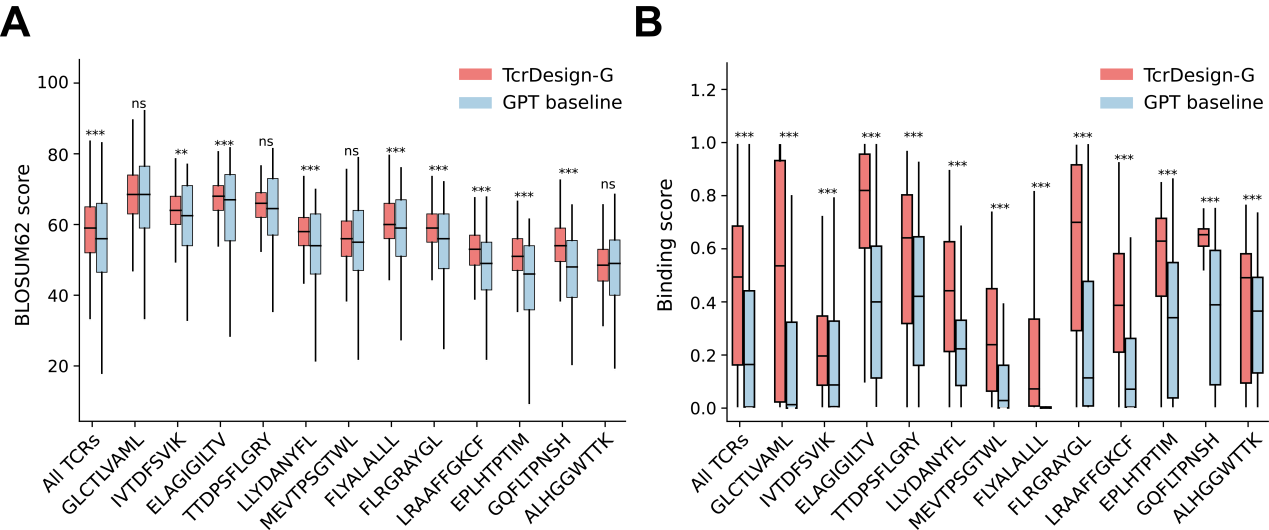

**Figure S11. The performance of TcrDesign-G and GPT baseline models was evaluated using beta chain information for representative epitopes.** The BLOSUM62 score (**A**) indicates the biological functional similarity between generated and natural βCDR3 sequences, while the binding score (**B**) reflects the predicted binding probability as assessed by TcrDesign-B. Two-sided Student’s t tests were applied (*p<0.05, **p<0.01, ***p<0.001, “ns” denotes “not significance”).

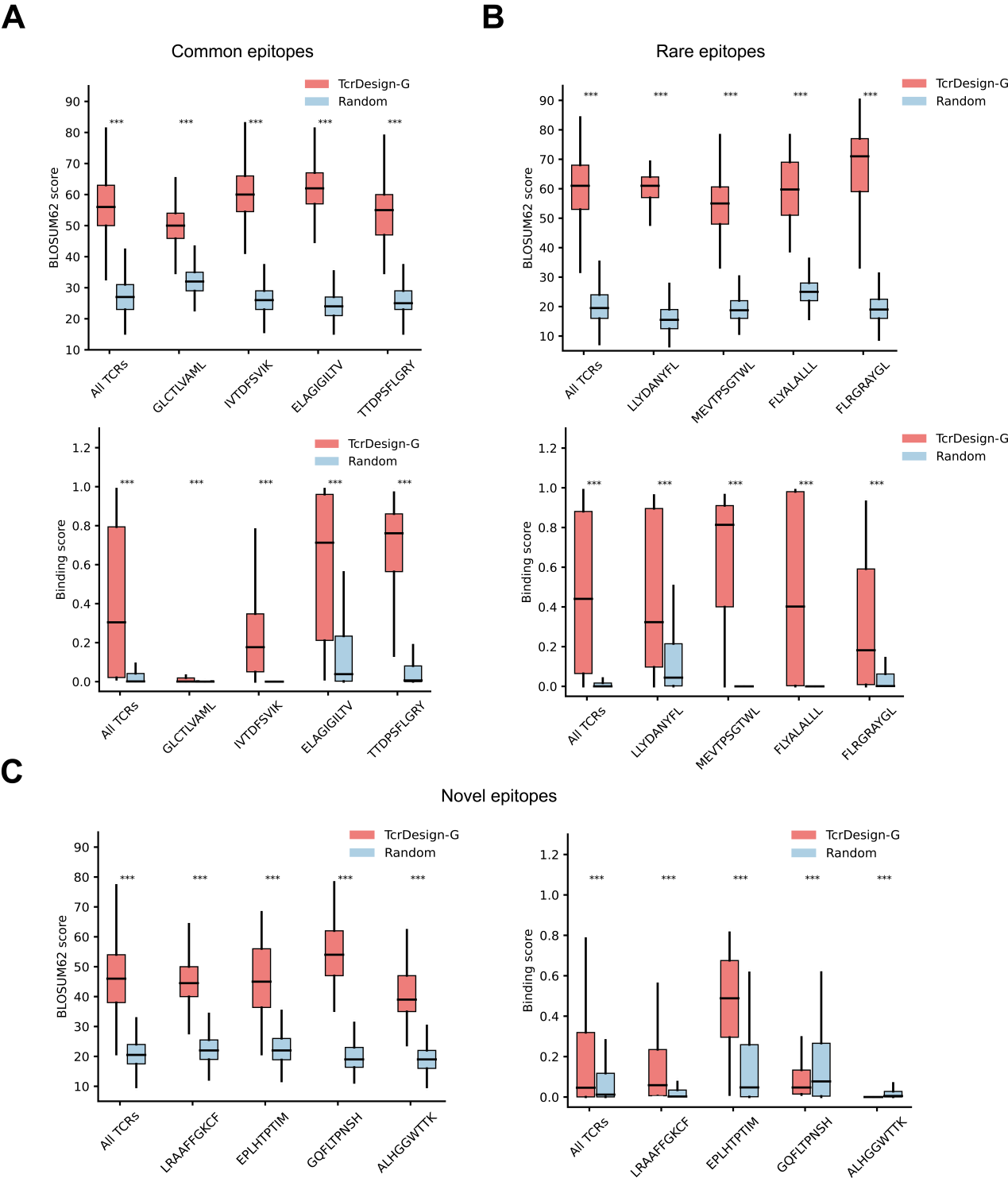

**Figure S12. The αCDR3s generated by TcrDesign-G showed high functional conservation compared to natural αCDR3s.** The performance of TcrDesign-G is evaluated using only alpha chain information for common epitopes (**A**) and rare (**B**) or novel (unseen) epitopes (**C**). The BLOSUM62 score indicates the biological functional similarity between generated and natural sequences, while the binding score reflects the predicted binding probability as assessed by TcrDesign-B. Two-sided Student’s t tests were applied (*p<0.05, **p<0.01, ***p<0.001, “ns” denotes “not significance”).

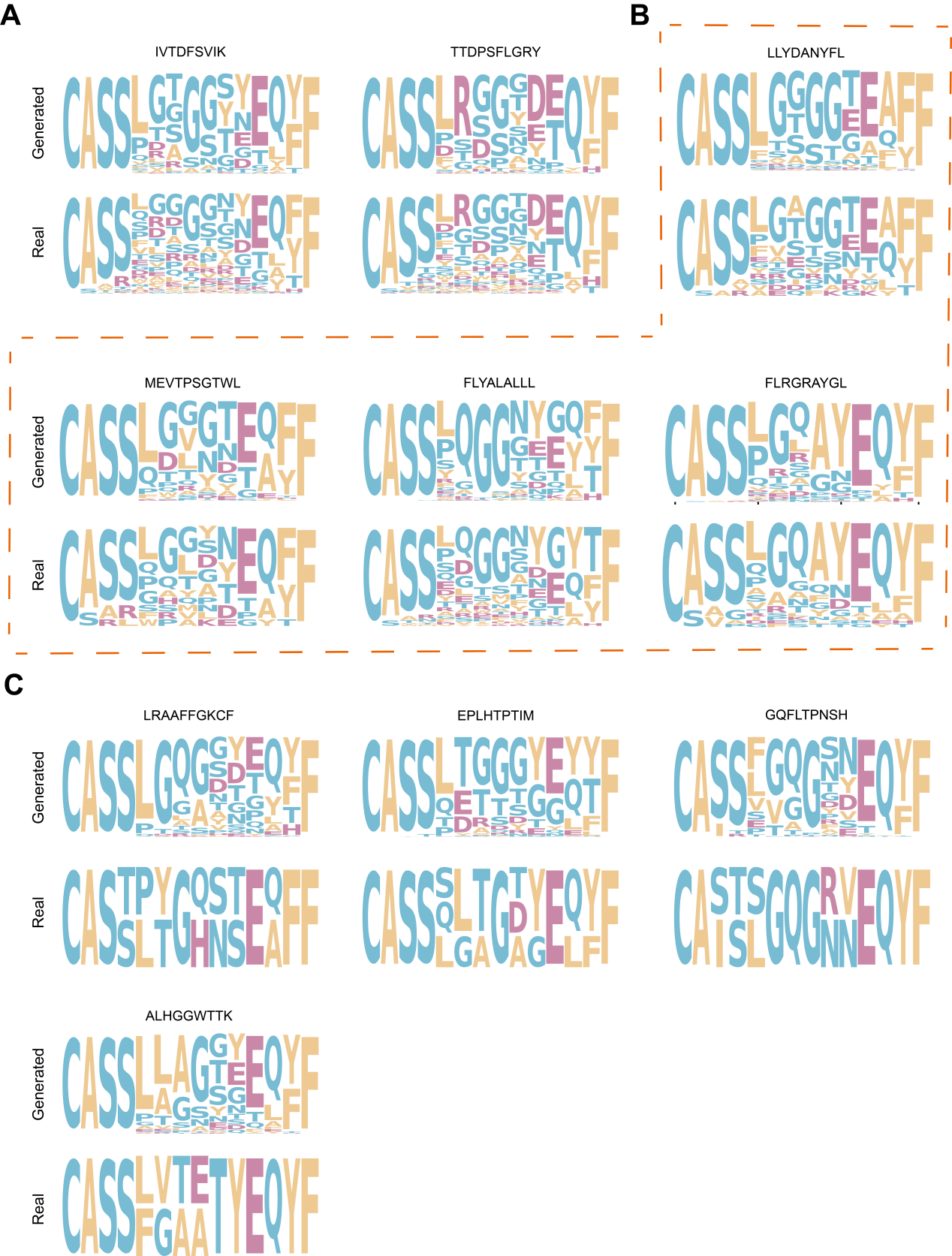

**Figure S13. Motif analysis for TcrDesign-generated βCDR3.** The results demonstrate the generation of 1,000 βCDR3 sequences for the common (upper), rare (middle) and novel (lower) epitopes, respectively, comparing these to the natural binding βCDR3 sequences.

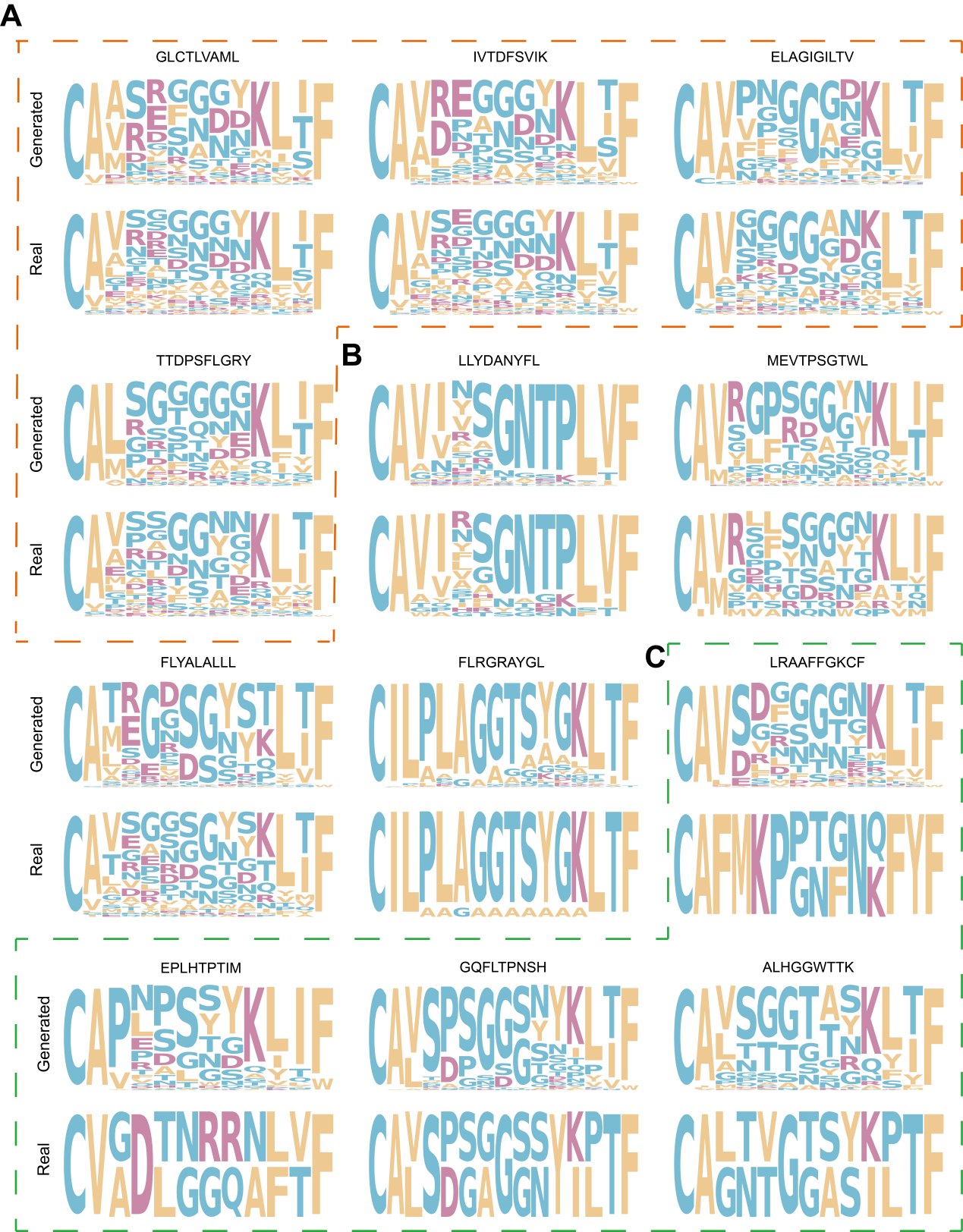

**Figure S14. Motif analysis for TcrDesign-generated αCDR3.** The results demonstrate the generation of 1,000 αCDR3 sequences for the common (upper), rare (middle) and novel (lower) epitopes, respectively, comparing these to the natural binding αCDR3 sequences.

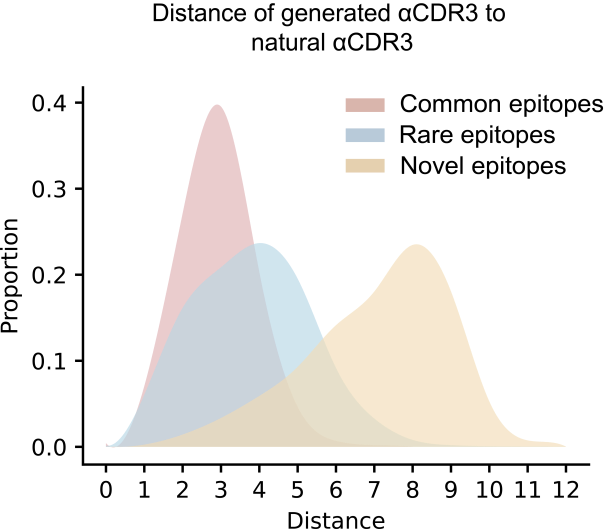

**Figure S15. Distance of generated αCDR3 to natural αCDR3.** The density plot illustrates the distribution of the minimum edit distances between 10,000 generated αCDR3 sequences for common, rare, and novel epitopes and their corresponding natural αCDR3 sequences.

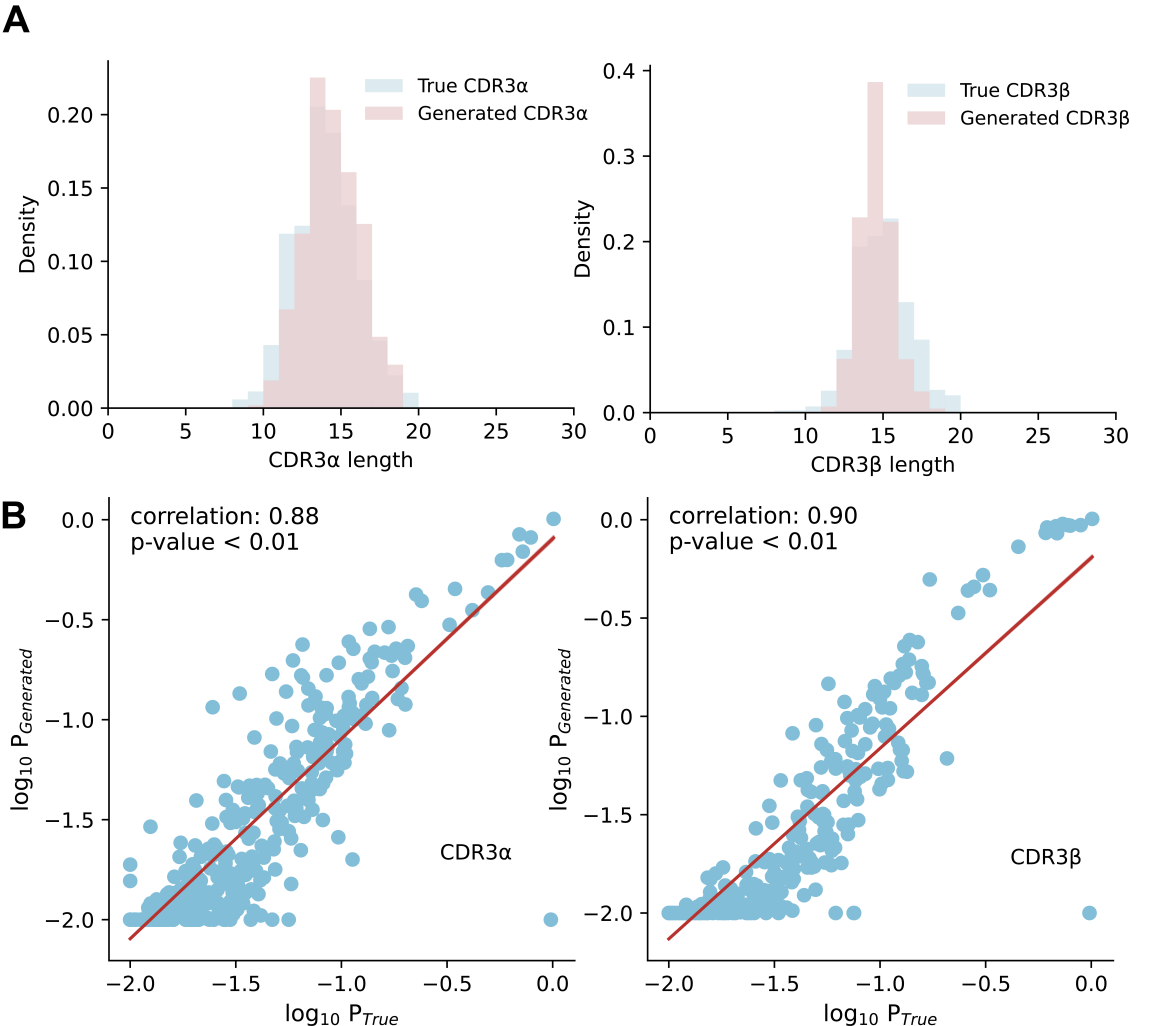

**Figure S16. Assessment of the biological plausibility of TcrDesign-generated TCRs.** (**A**) Density distributions of CDR3α (left) and CDR3β (right) loop lengths. The generated TCRs (pink) closely match the length profile of natural TCR repertoires (light blue). (**B**) Scatter plots comparing the log-probabilities (the log probability distribution of the 20 standard amino acids at each position) of generated versus true TCR sequences for CDR3α (left, r = 0.88) and CDR3β (right, r = 0.90). High correlation (p < 0.01) along the y = x line (red) indicates that TcrDesign produces sequences with statistical properties consistent with natural TCRs.

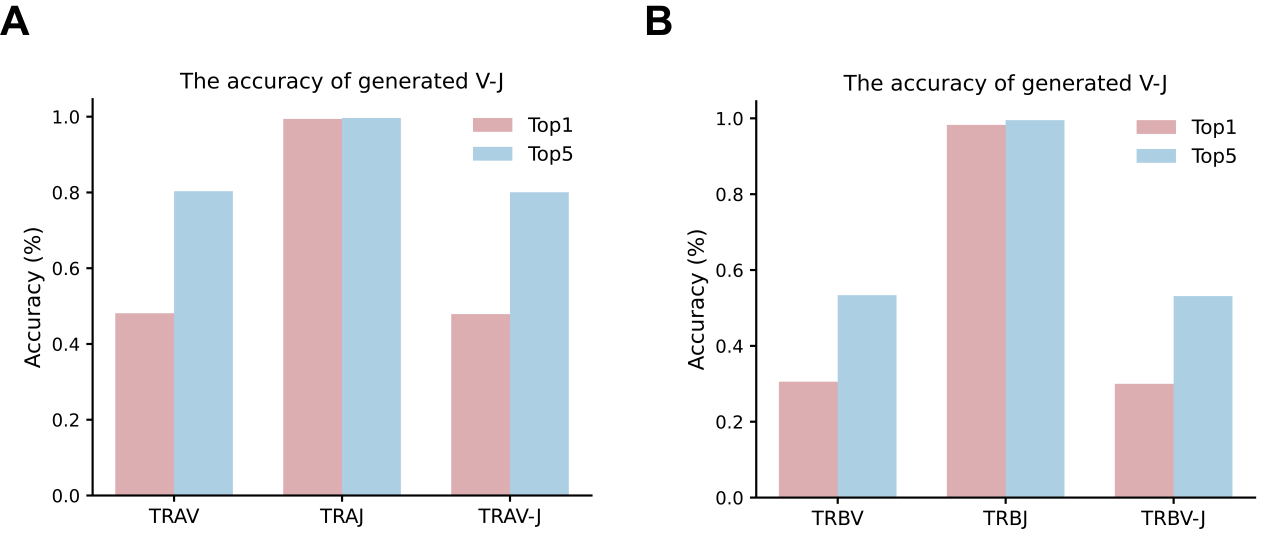

**Figure S17. Inference VJ genes from CDR3 sequences.** (**A**) Accuracy of generated VJ genes for αCDR3s from top1 or top5 predictions. (**B**) Accuracy of generated VJ genes for βCDR3 from top1 or top5 predictions.

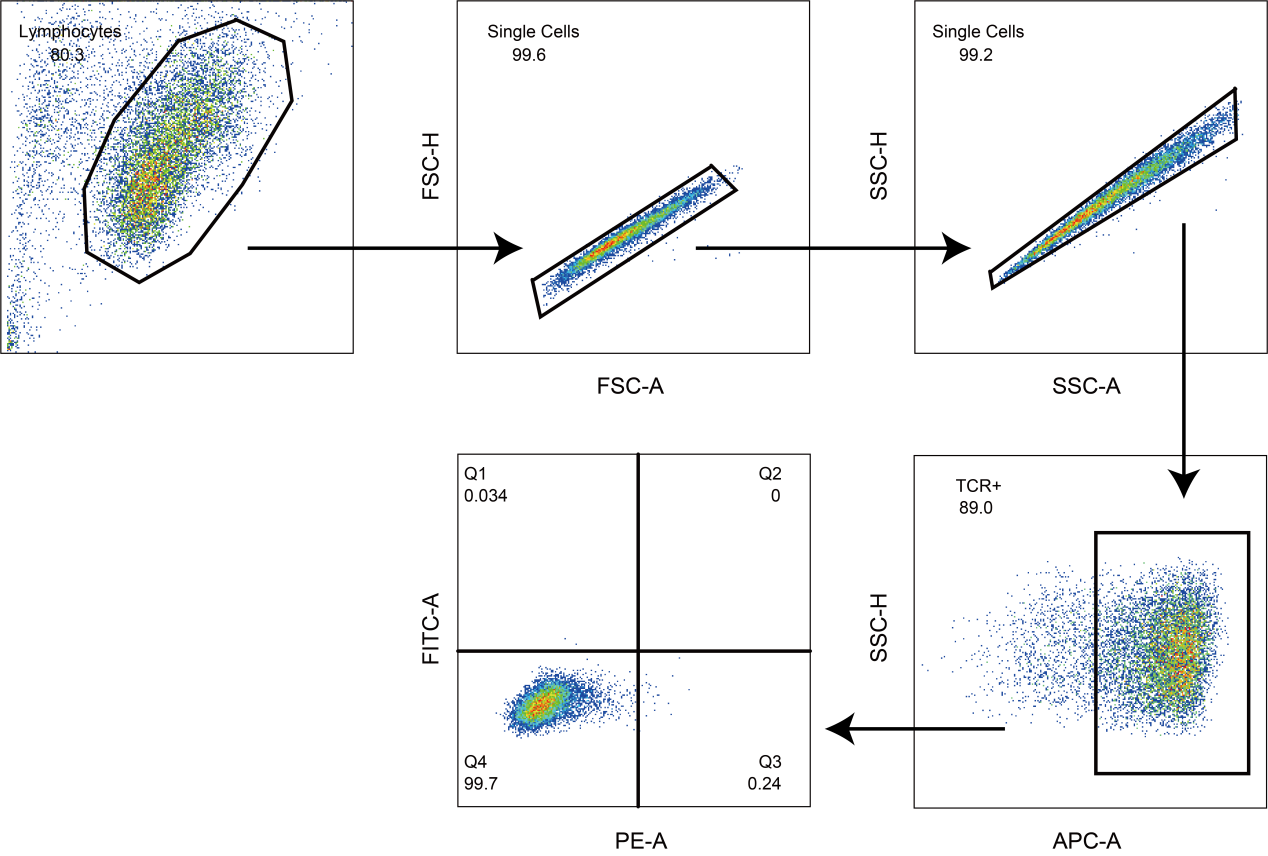

**Figure S18. Evaluation of the frequency of CD69-positive and ZsGreen-positive Jurkat T cells by flow cytometry.** Gating strategy and assessment of the TCR-Engineered Jurkat T cells after co-incubating with target cells, staining by APC anti-human TCR α/β antibody and PE anti-human CD69 antibody, negative control sample is shown here.

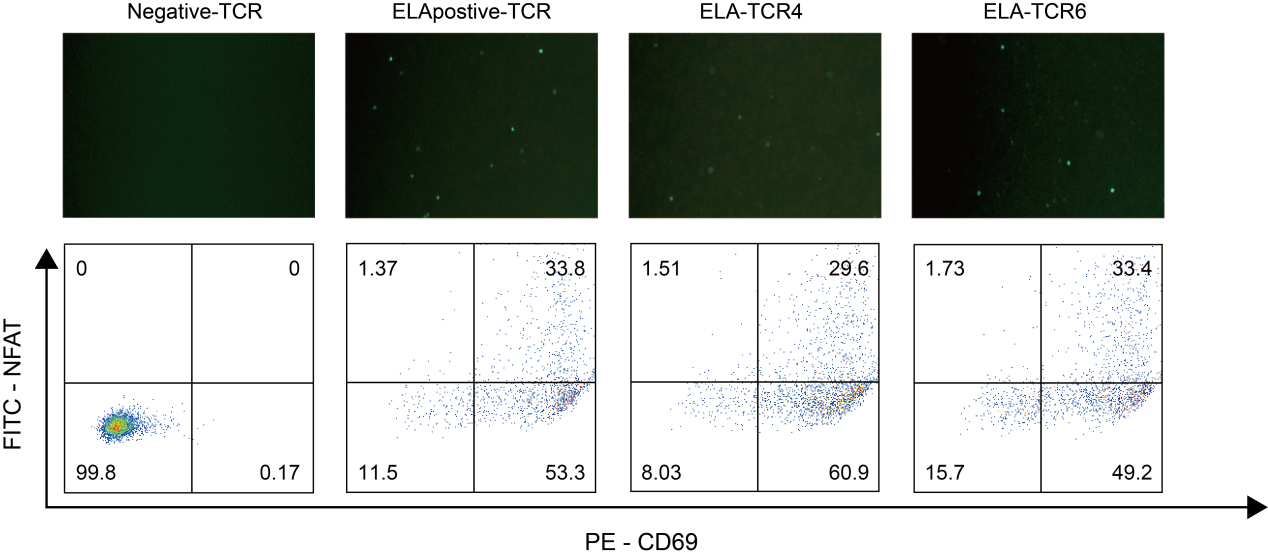

**Figure S19. Fluorescence imaging and flow cytometory analysis of engineered Jurkat T cells.** Jurkat NFAT-ZsGreen reporter cells, expressing candidate TCRs, were co-cultured with ELAGIGILTV-pulsed T2 cells. After 24 hours, flow cytometry and fluorescence imaging were performed. ELA-TCR4 and ELA-TCR6 showed significant signals for CD69 and NFAT-ZsGreen in contrast to negative control.

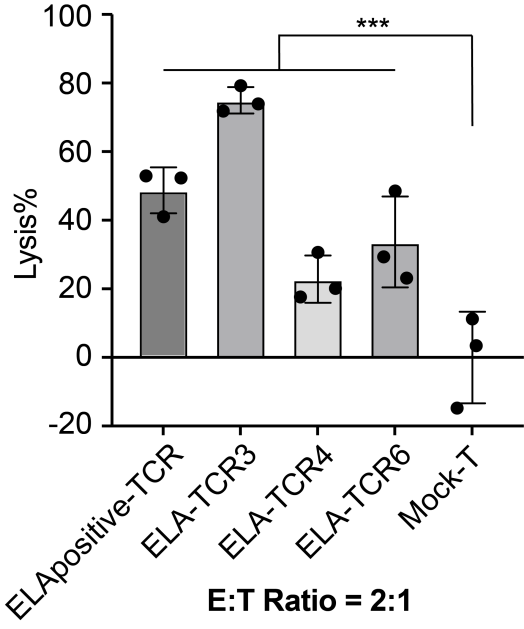

**Figure S20. Lysis percentage of ELAGIGILTV pulsed T2-luciferase cells.** Mock T cell and TCR-engineered T cells (ELA-positive, ELA3, ELA4, ELA6) were co-cultured with T2-luciferase cells pulsed with 1 μg/mL ELAGIGILTV peptide at an effector-to-target (E:T) ratio of 2:1 for 24 hours. Target cell lysis was quantified using a bioluminescence-based cytotoxicity assay. TCR-engineered T cells exhibited significantly enhanced antigen-specific cytotoxicity compared to Mock T cells (*p < 0.001, one-way ANOVA with Dunnett’s post-test). Data represent mean ± SD of triplicate experiments.

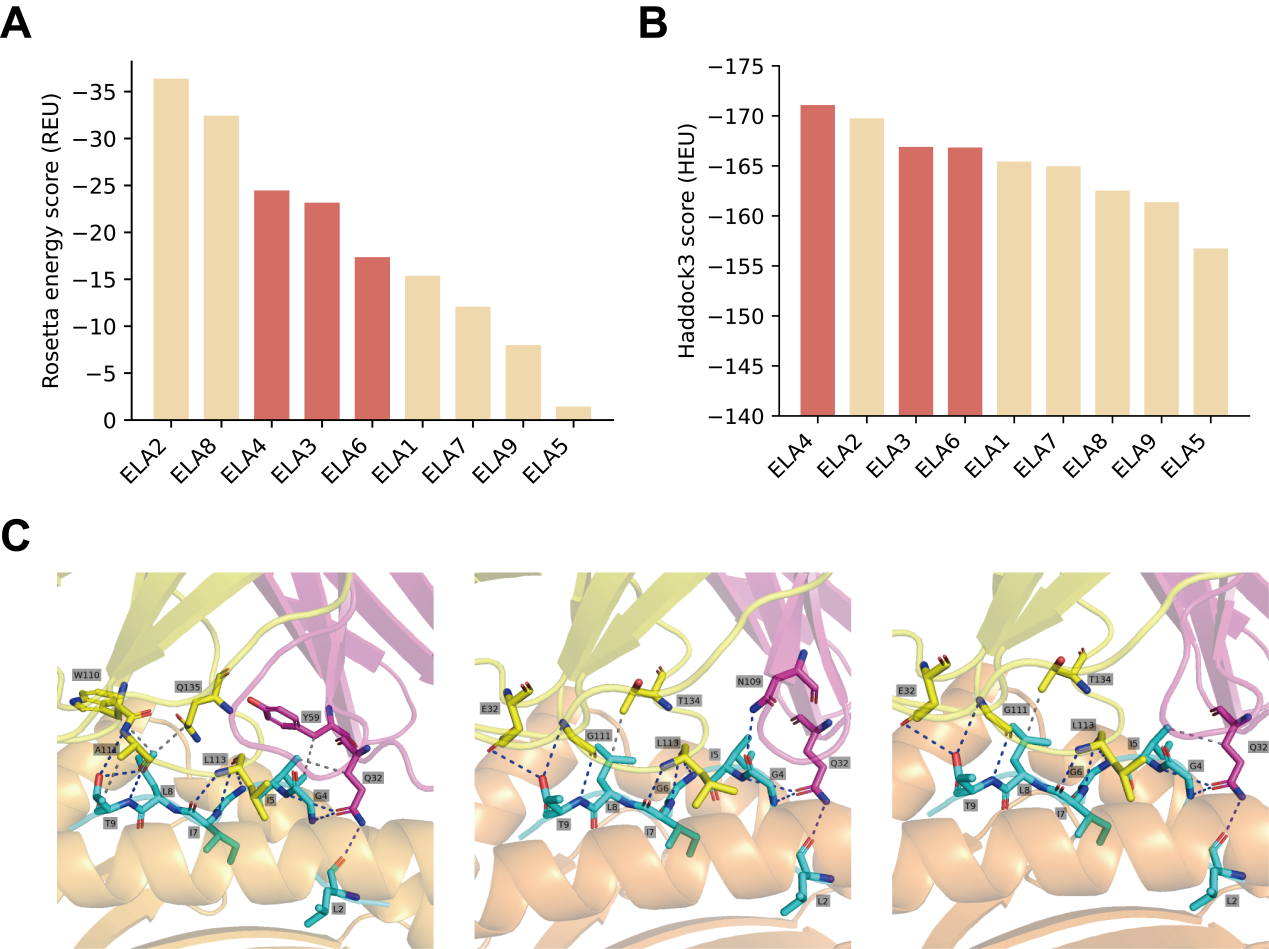

**Figure S21. Computational analysis of binding energy and interaction patterns for TCR–pMHC complexes.** (**A**) Rosetta energy scores (REU) for TCR-pMHC complexes. Lower values indicate higher structural stability. Functionally validated TCRs (ELA3, ELA4, ELA6) are highlighted in red and exhibit among the lowest energy scores. (**B**) Haddock3 scores (HEU) reflecting TCR-pMHC docking affinity. Lower scores correspond to stronger binding. The same validated TCRs (highlighted in red) consistently rank near the top. (**C**) Predicted binding interfaces for ELA3 (left), ELA4 (middle), and ELA6 (right). TCR α and β chains are shown in magenta and light yellow, respectively; MHC chain in orange; the antigenic peptide (ELAGIGILTV) is represented as a cyan stick model. Hydrogen bonds and hydrophobic interactions are indicated with blue and gray dashed lines, respectively.

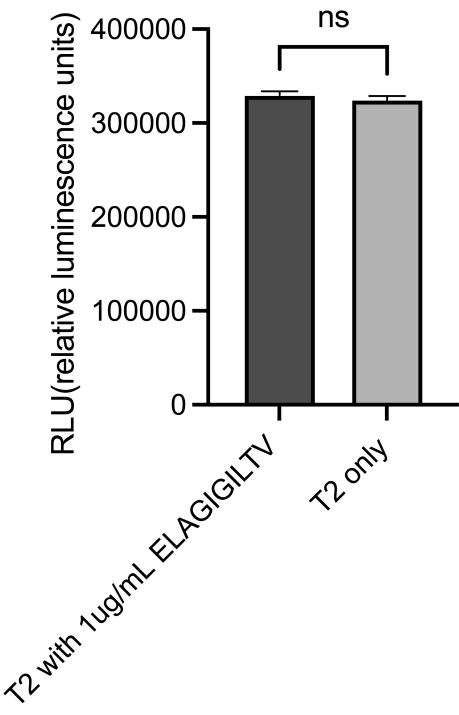

**Figure S22. Baseline luminescence of T2-NFAT-luciferase reporter cells is unaffected by peptide pulsing.** T2 cells stably expressing an NFAT-driven luciferase reporter were incubated with 1μg/mL ELAGIGILTV peptide or medium alone (T2 only) for 16 hours. Luminescence was measured in relative luminescence units (RLU). No significant difference (ns, p > 0.05) was observed between the peptide-pulsed and unpulsed groups, confirming that peptide incubation at this concentration does not alter baseline cell viability.

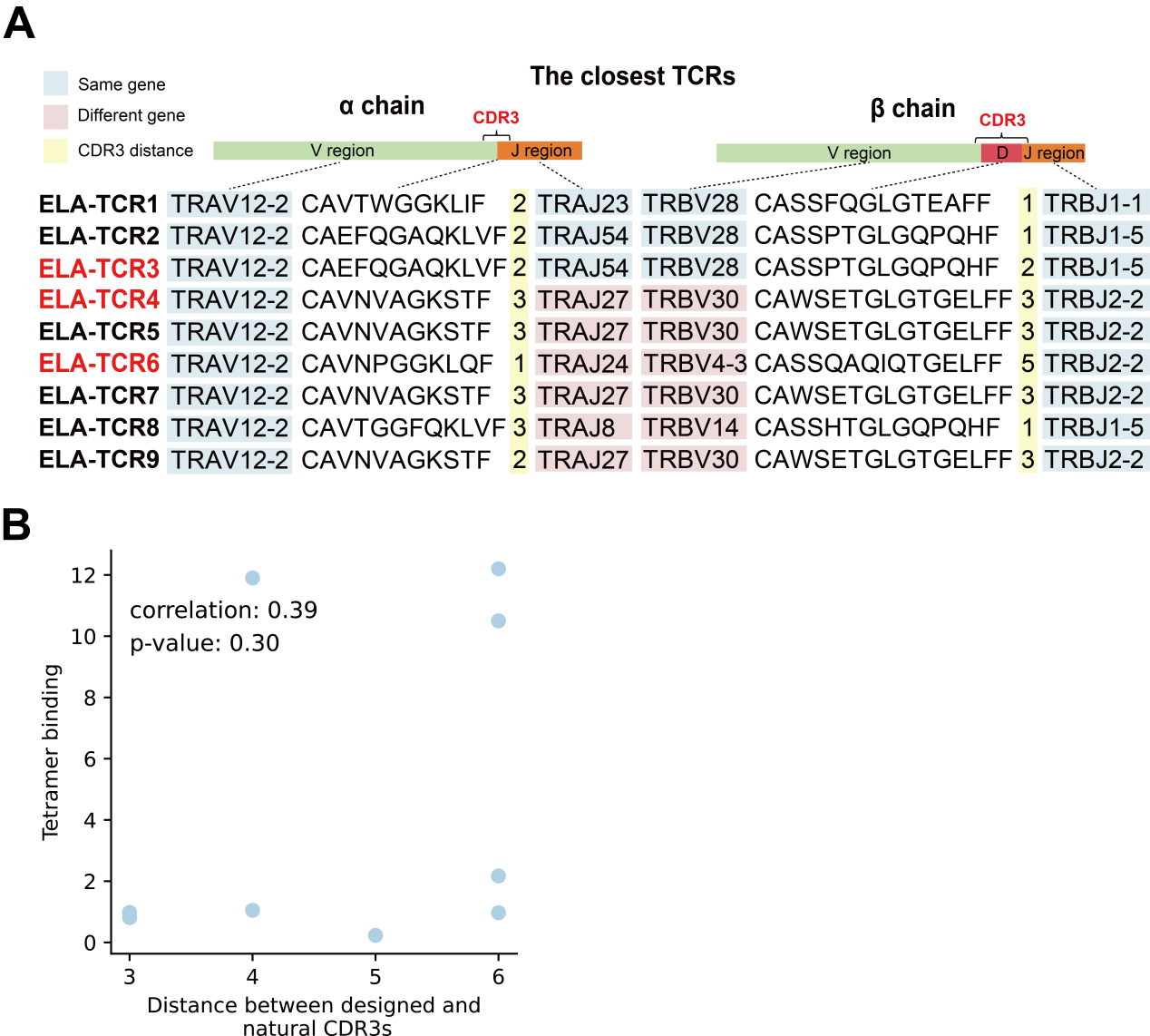

**Figure S23. Computational analysis of designed TCRs targeting ELAGIGILTV and their closest natural TCRs.** (**A**) Comparison of 9 designed TCRs with their closest ELAGIGILTV-specific natural TCRs. Light blue indicates identical gene usage between the designed and natural TCRs, while pink indicates different gene usage. The highlighted yellow column represents the edit distance between the closest natural CDR3 (left) and the designed CDR3. ELA-TCR3, 4, and 6, highlighted in red, denote experimentally reactive TCRs. (**B**) Scatter plot illustrating the correlation between the tetramer-binding fraction and the edit distance between natural and designed CDR3 sequences.

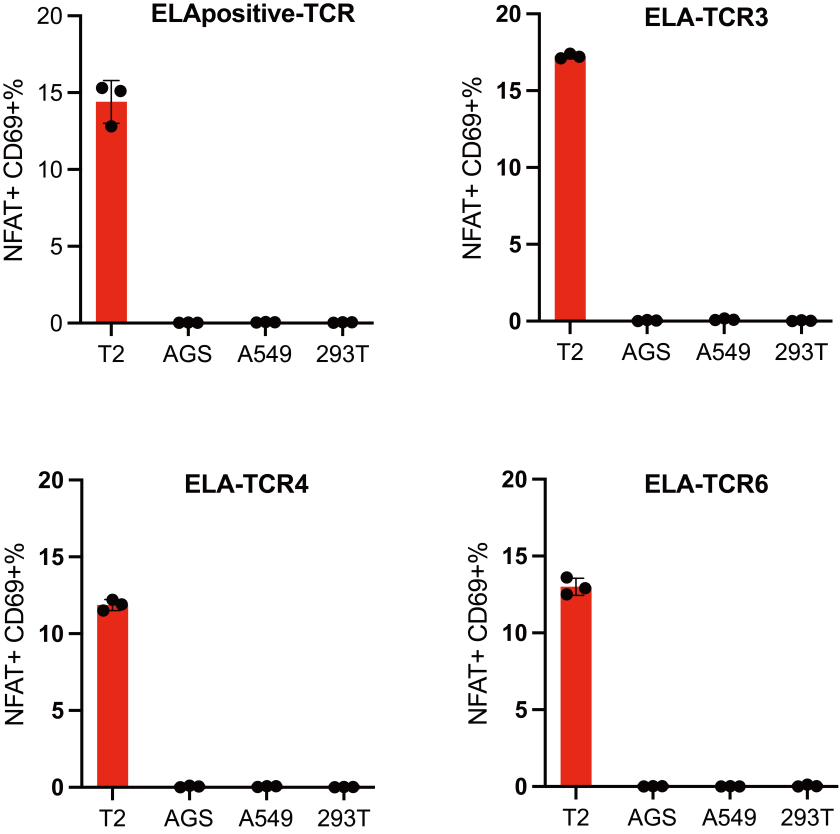

**Figure S24. Assessment of off-target reactivity for designed TCRs.** NFAT-luciferase reporter assay was performed to evaluate T cell activation (quantified as NFAT+ CD69+ percentage) induced by TCRs specific to the ELAGIGILTV epitope across multiple human cancer cell lines. T2 cells pulsed with the target peptide served as a positive control, while AGS, A549, and 293T cell lines-representing diverse endogenous peptide backgrounds-were used to assess off-target activation. ELA-TCR3, ELA-TCR4, ELA-TCR6 showed specific activation only on peptide-loaded T2 cells, with minimal reactivity against other cell lines, demonstrating low risk of off-target effects.

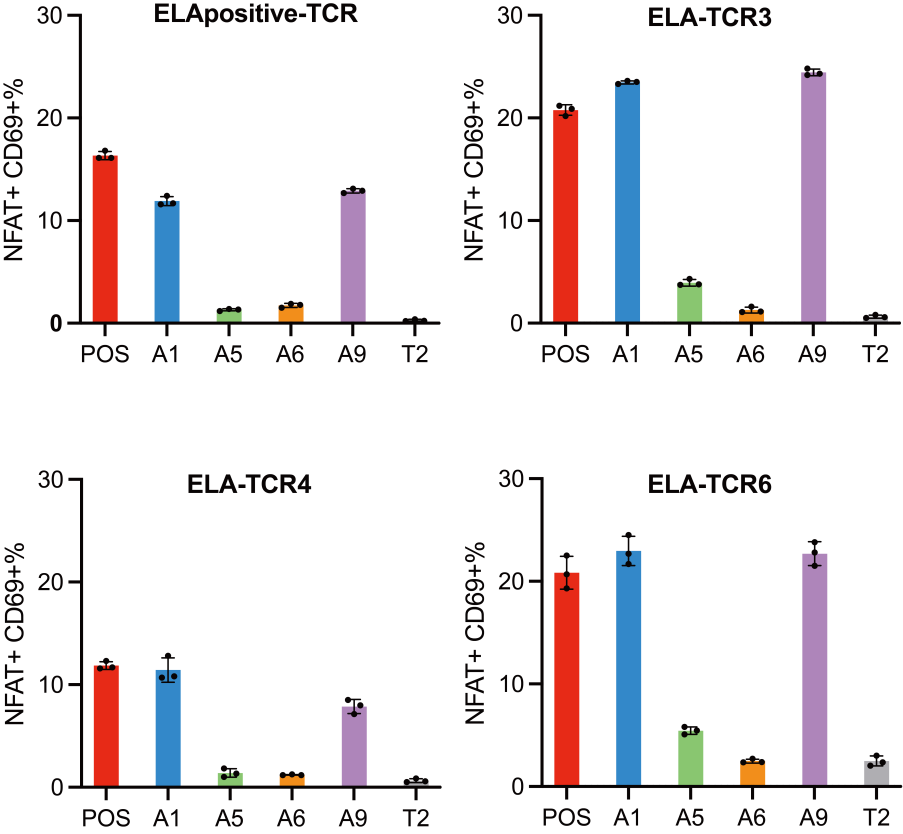

**Figure S25. Evaluation of off-target reactivity for designed TCRs using alanine scanning mutagenesis.** Alanine scanning was conducted on the ELAGIGILTV peptide, with candidate peptides predicted to bind HLA-A*02:01 selected using netMHCpan v4.1. Off-target effects of similar peptides were assessed via co-incubation of Jurkat cells expressing the indicated TCRs (ELA-TCR, ELA-TCR3, ELA-TCR4, or ELA-TCR6) with T2 cells pulsed with the respective peptides. T cell activation was quantified as the percentage of NFAT+ CD69+ cells. POS: pulsed with the ELAGIGILTV peptide; A1, A5, A6, A9: pulsed with the peptide which alanine substitutions at peptide positions 1, 5, 6, and 9, respectively; T2: T2 cells only, without peptide pulsing.
